# Neural manifolds that orchestrate walking and stopping

**DOI:** 10.1101/2025.11.08.687367

**Authors:** Salif Komi, Jaspreet Kaur, August Winther, Madelaine C. Adamsson Bonfils, Grace A. Houser, R.J.F. Sørensen, Guanghui Li, Karen Sobriel, Rune W. Berg

## Abstract

Walking, stopping and maintaining posture are essential motor behaviors, yet the underlying neural processes remain poorly understood. Here, we investigate neural activity behind locomotion and its walk-to-stop transition. Based on a new theory of the lumbar spinal cord^1, 2^ we propose and predict that spinal population activity contains limit cycle dynamics to drive walking and fixed-point attractors for stopping. To test these predictions we record neural activity in lumbar cord of freely moving rats using Neuropixels probes^3^. To control stopping, we stimulate a brainstem nucleus, known to induce motor arrest^4–7^. We find: During locomotion, the population activity of lumbar spinal neurons exhibits rotational dynamics^8–10^. These dynamics unfold within a low-dimensional locomotor manifold^11, 12^, a looping set of trajectories that serves as the repeating signature of locomotion, that also behaves as a limit-cycle attractor. Shortly before stopping, the neural state rapidly changes from the locomotor manifold to a ‘postural’ fixed point attractor. When kicking the state out of the fixed point using perturbations it shifts to a nearby albeit different fixed point. Repeated stoppings form a local quasi-continuum of fixed points representing various poses – i.e. a postural manifold. These observations are in agreement with our theory, which further indicates the mechanistic roles for subpopulations of spinal interneurons for controlling walking and stopping. Besides explaining the data, our theory makes further predictions to be tested in future experiments.

## Main

How does a cat gracefully walk and suddenly freeze when it spots a mouse? Walking is essential^13,14^, but equally important is the ability to stop the ongoing movement, for example, in the sudden appearance of an obstacle, a threat, or simply when needing a rest. This seamless coordination between movement and posture has long fascinated physiologists; Sherrington famously remarked that ‘posture follows movement like a shadow’^15^ illustrating the idea that stability and motion are dynamically intertwined parts of a single motor process. Although Sherrington’s observation conceptually linked movement and posture, the underlying neural mechanisms that coordinate instant switching between movement and immobility are largely unaddressed. Conventional central pattern generator (CPG) models and experiments have aimed to explain the basis of locomotion with tremendous contribution^15–21^, but few if any explain the mechanics of stopping and maintaining posture. The experiments that have shaped conventional wisdom are based on models in which brain-spinal communication is severed, and hence motor transitions are absent. The experiment necessary to address walk-to-stop is in awake animals performing volitional motor behaviors where brain-spinal communication is intact, but these have been rare for technical reasons until now^22–24^.

The recording of spinal neurons during freely moving animals is important for another reason. Recent studies revealed rotational population activity where all phases of the step cycle are represented^10,25,26^, which conflicts with the classical CPG models that have modular alternation in two-phases. This highlights the mystery of walk-to-stop: How can a biphasic system be stopped and kept in any phase when it only has two phases? We propose that understanding the issue of walk-to-stop is the key to understanding walking itself. A natural starting point would be a network model that has rotational population dynamics and then explore potential stopping-mechanisms. Recently,

### Walking and stopping: Theory

In our theory, the spinal circuitry can generate rhythms and patterns purely by virtue of spatial organization without relying on rhythmogenic cellular properties^1^. The network consists of cell types that are represented by their longitudinal projections in the mouse lumbar cord (**Fig. 1a**). Critically, the total projection pattern of excitation and inhibition (E/I), the ‘projectome’, needs to take the shape of a ‘Mexican hat’, that is, with recurrent local excitation balanced by long-range inhibition along the rostro-caudal axis^27^ (**Fig. 1b**). This architecture is inspired by the ‘Ring network’, which is seen elsewhere in the nervous system, such as for head-direction circuits where a ‘bump’ of activity can remain stable due to the continuous attractor properties of the network^28,29^. In these networks, the stability depends on the symmetry of the E/I projections, and if symmetry is broken, the bump will move. If we imagine the spinal network having a similar Mexican hat projectome with asymmetry, a bump of activity that travels could serve as a sequence generator and recruit muscle groups. The recruitment occurs as the bump passes and activates the pool of motoneurons (MN). Since MN pools are organized with a spatial shift of various flexor/extensor pools, propagation will induce a sequence of muscle contractions associated with normal walk^30,31^ (**Extended Data Fig. 1**). The bump itself emerges at the rostral end of the spinal cord due to local instability through recurrent excitation, which is activated by descending drive. As the bump propagates far enough downwards it allows a new bump to emerge, hence the network is effectively both a rhythm- and pattern-generator purely due to the network architecture.

**Fig. 1.**
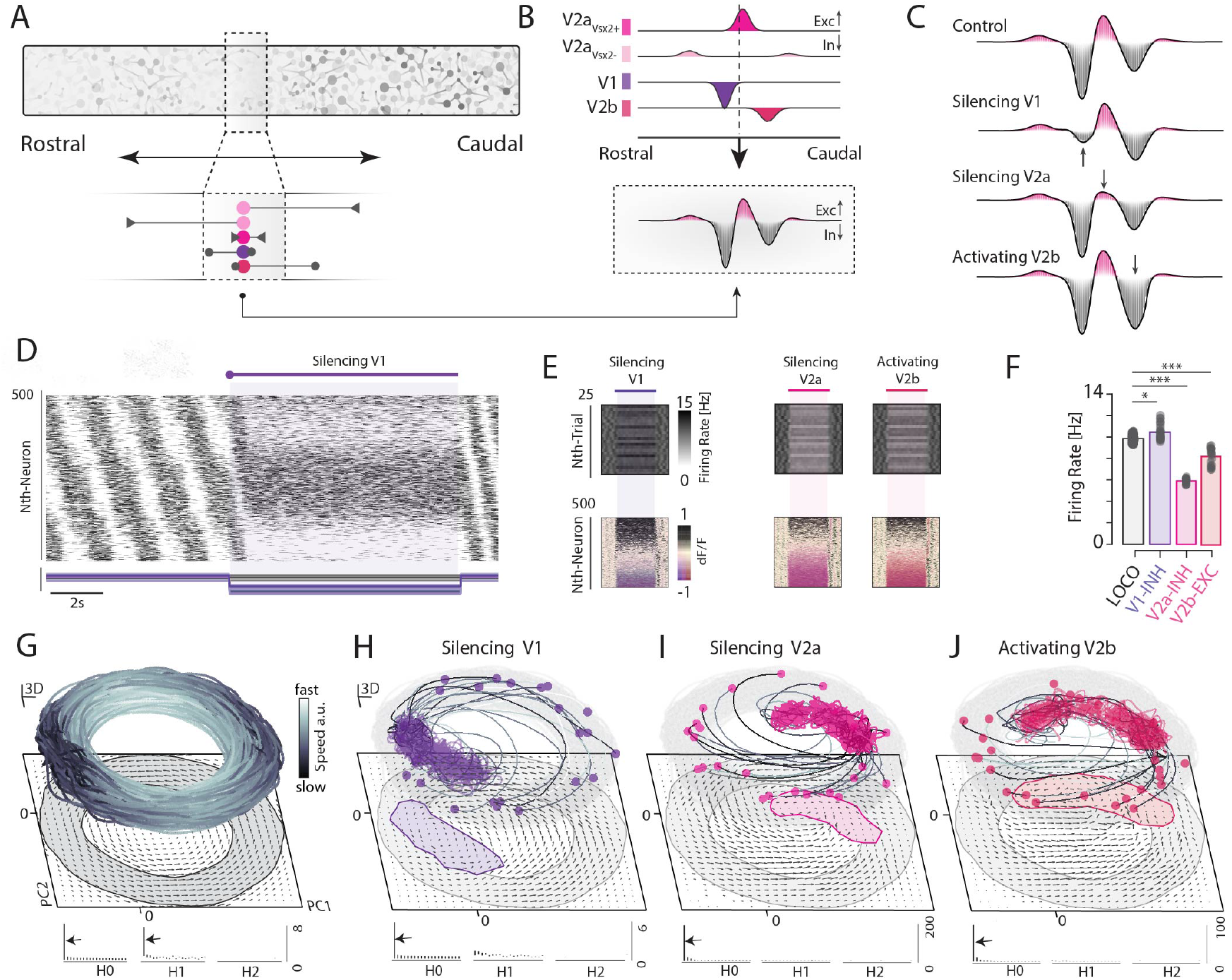
Model of spinal motor network and the walk-to-stop transitions: Bifurcation from limit cycle to a fixed point attractor. **a**, Projections of spinal interneurons from a given point in the longitudinal direction, are composed of various cell types (colors). **b**, Local recurrent excitation, (V2a-Vsx2+, magenta), long-range excitation (V2a-Vsx2-, pink), ascending inhibition (V1, purple), and descending inhibition (V2b, red), together form a Mexican hat-shaped total input versus distance, i.e. projectome (bottom). **c**, illustration of the projectome shape during cell-specific gain-modulation. Asymmetry is embedded from the start (control, top). Silencing V1, V2a-Vsx2+, and activating V2b, below respectively (arrows). **d**, Population activity in the model show sequential progression and stopping when silencing V1 (gain put to zero, bottom, purple). **e**, Population firing rates (gray scale, top), and the sorted according to change (color map, bottom), when manipulating the three cell types. **f**, Mean population firing rates of locomtion (left), and the three manipulations. **g**, State space activity during walking, has rotation with a solid ring structure as assessed by homology (arrows, bottom). **h**, When silencing V1, activity bifurcates to a new state, the postural manifold (purple). The manifold is topologically equivalent to a solid (only one line in persistent homology, bottom). **i-j**, similar bifurcations during the manipulations of V2a-Vsx2+ and V2b. we developed a theory based on anatomical principles that offers clear predictions for locomotion, rotational dynamics, and cessation of movement while maintaining muscle tone^1,10^. Here, we first outline these predictions and then test them in experiments by recording the activity of lumbar spinal neurons in freely moving rats using Neuropixels probes.

Now, a walk-to-stop transition involves stopping the bump at a location that corresponds to a particular posture. Recurrent local excitation will maintain the bump and support muscle tone. To test this idea, we focus on ipsilaterally projecting neurons, as it is known that commissural connections are not required for rhythm generation. Key candidates are the inhibitory V1 and V2b cells, with asymmetric ascending and descending projections^32–34^, together with local excitatory V2a cells^21,35^, although others may also contribute. By modeling such a network, we find that inhibiting V1 or V2a, or activation of V2b, has contrasting effects on the projectome (**Fig. 1c**). However, these manipulations induce similar motor arrest, as characterized by a transition from rhythmic to tonic firing (**Fig. 1d-j**). When V1 neurons are sufficiently inhibited the propagation of the bump stops altogether (**Fig. 1d**). The mean firing rate also changes, some neurons increase, others decrease (**Fig. 1e-f**).

Analyzing the population activity in state space using principal component analysis (PCA) of the firing rates, a low-dimensional ring-shaped manifold appears, which we call the ‘locomotor manifold’ (**Fig. 1g**). The manifold is topologically a solid ring, rather than a torus, as revealed by the specific homology barcodes^28^ (*H*_0_ and *H*_1_, bottom **Fig. 1g**). Since the trajectories remain stable inside the manifold, the behavior resembles a limit cycle attractor. Although traditional limit cycles are circles with no thickness, the thickness here comes from slight changes in parameters, e.g. the descending drive or the neuronal gain. Hence, the manifold contains a set of nested limit cycles that arise from changes in the parameters. For example, the radius of rotation would increase as more muscular force is needed for a particular step^10^. When inducing a stop by modifying the effective projectome, the state goes from the periodic orbit to a stable fixed point. Since the limit cycle orbit of a particular set of parameters has an associated fixed point, the set of limit cycles has a corresponding set of fixed points, which form a quasi-continuous region called the ‘postural manifold’. (**Fig. 1h-j**). Upon initiation of the stop, trajectories converge from the locomotor manifold onto the postural manifold. The postural manifold is solid (barcode in *H*_0_, bottom). Since the postural manifold is not ring-shaped, it cannot support a static pose in all phases of the walking cycle. In summary, this theory of movement generation predicts two types of attractor manifolds for spinal network activity, and during the walk-to-stop neural state, it can make a quick transition from the locomotor to the postural manifold.

### Neural activity during walking

To verify the theory, we chronically implant high-density Neuropixels probes^3,22,36^ into the lumbar spinal cord of rats (**Methods**). The spinal probe is implanted at an angle of 30-40° in the lumbar region (L1-L5) (**Fig. 2a**). To better control stopping, we optogenetically activate a nucleus known to induce arrest, the pedunculopontine nucleus (PPN)^4–7^. The probe locations are reconstructed using histology and imaging (**Fig. 2b** and **Extended Data Fig. 2-3**). The kinematics of the animal walking on a speed-adaptive treadmill^37^ is extracted using videography (**Extended Data Fig. 4, Supplementary Video 1**). Although movement artifacts are a general challenge, their impact can be reduced by clustering spikes and following units across sites using a drift correction technique^36^ (**Fig. 2c** and **Extended Data Fig. 5**). Hence, there are periods of stable recording that allow for the investigation.

**Fig. 2.**
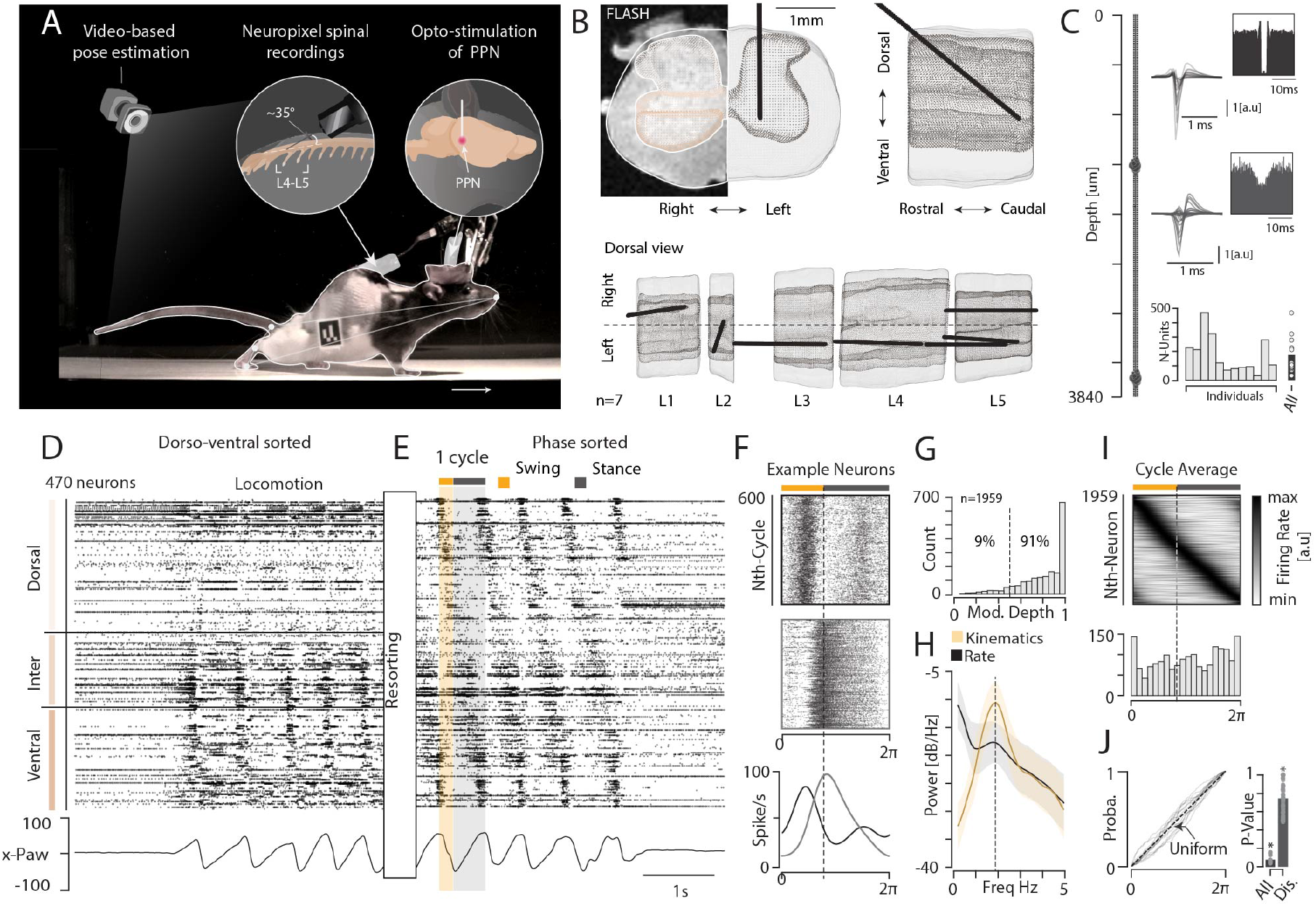
Lumbar cord has rotational population dynamics during walking. **a**, Experimental setup: spinal activity is recorded using an implanted Neuropixels probe (left) and optogenetics in the brainstem nucleus, PPN (right). Animal walks freely on adaptive treadmill and kinematics is extracted using videography. **b**, Reconstruction of probe locations in lumbar cord segments (L1-L5, n=7/12 animals shown) using MRI. **c**, Sample units on probe, top is a single-unit (‘good’, multiple channels plotted together) with a auto-correlogram value of zero at origin (inset); bottom a multiunit (not good) with non-zero value at origin (inset). Number of good units across animals, average ∼160 (bottom). **d**, Raster of spiking (n=440, sample R173) localized dorsal(top)-ventral(bottom). Bottom: kinematics. **e**, continuation of raster with phase-sorted units. Stance/swing phase indicated (orange/grey). **f**, step-cycle normalized rasters for two sample neurons over repeated step cycles (y-axis), averaged shown (bottom). **g**, modulation depth distribution across population, indicates majority is rhythmic (91%). **i**, Pooling phase-sorted data across animals show continuous cycle progression (top), phase distribution (middle). **j**, Cumulative phase distributions (bottom) indicate near-uniform distribution (linear increase), albeit distinguishable from linear (KS2-test, bottom right, n=12 animals with N=1956 neurons, p-values indicated).

Spiking neurons are found along the dorso-ventral axis and many had rhythmic activity during walking (**Fig. 2d**). Individual cells have preferred firing phases during the walking cycle, and when the neuronal order is arranged according to the phase, a distinct firing sequence appears (**Fig. 2e**). The sequence repeats for every step and the phase-preference remains across cycles (**Fig. 2f**). Most neurons are rhythmic (91%, **Fig. 2g**) and their frequencies aligns with the kinematics (**Fig. 2h, Extended Data Fig. 6**). When combining all neurons across trials and animals (n=12 animals), the sequence has a near-uniform distribution (**Fig. 2i**). This phenomenon, known as rotational population dynamics, has previously been characterized in other motor behaviors and elsewhere in the nervous system^8, 10, 38^.

### Neural transition behind walk-to-stop

Having established rotational population activity during walking, we next examine the activity associated with volitional stopping. During such a transition, the firing shifts from sequential to tonic activity as the animal assumes its posture (**Fig. 3a**). During walking, population activity in state space (PCA)^11^ enters a low-dimensional manifold, the locomotor manifold – where only three principal components of the averaged firing rates across step cycles are needed to explain most of the variance (>80%, **Fig. 3b, Extended Data Fig. 7**). This is a well-defined circular manifold, where the onsets of the swing and the stance are confined to specific locations (dots, **Fig. 3c**). The motion slows down as the leg enters the stance phase and achieves mechanical stability, and speeds up as it starts the swing phase (gray scale, **Fig. 3d**). The motion in and outside the manifold can be visualized using flow fields, with arrows indicating the change in the state of the network (**Fig. 3e**). As in the model (**Fig. 1g**) the manifold is topologically a solid ring (barcodes indicated, bottom). The flow inside the locomotor manifold is well organized (large ‘alignment’, left, **Fig. 3g**) and with greater flow than outside (**Fig. 3h**), indicating a set of limit-cycle attractors. These features of the locomotor manifold are in agreement with the predictions of the theory. During voluntary stops, the state converges to a solid region – the ‘postural manifold’ – composed of multiple fixed-points (**Fig. 3f**). Contrary to the locomotor manifold, the flow inside the postural manifold is disorganized, i.e. small alignment and flow indicating that movement is only fluctuating around stable fixed-points (**Fig. 3g-h**). These observations along with the alignment of behavioral parameters indicate that walking and its transition to stopping involve a bifurcation from a limit-cycle orbit to stable fixed-point states. As in theory, the postural manifold is not ring-shaped and therefore does not support poses for all phases of the cycle, which could be due to network properties and stability constraints of the biomechanics.

**Fig. 3.**
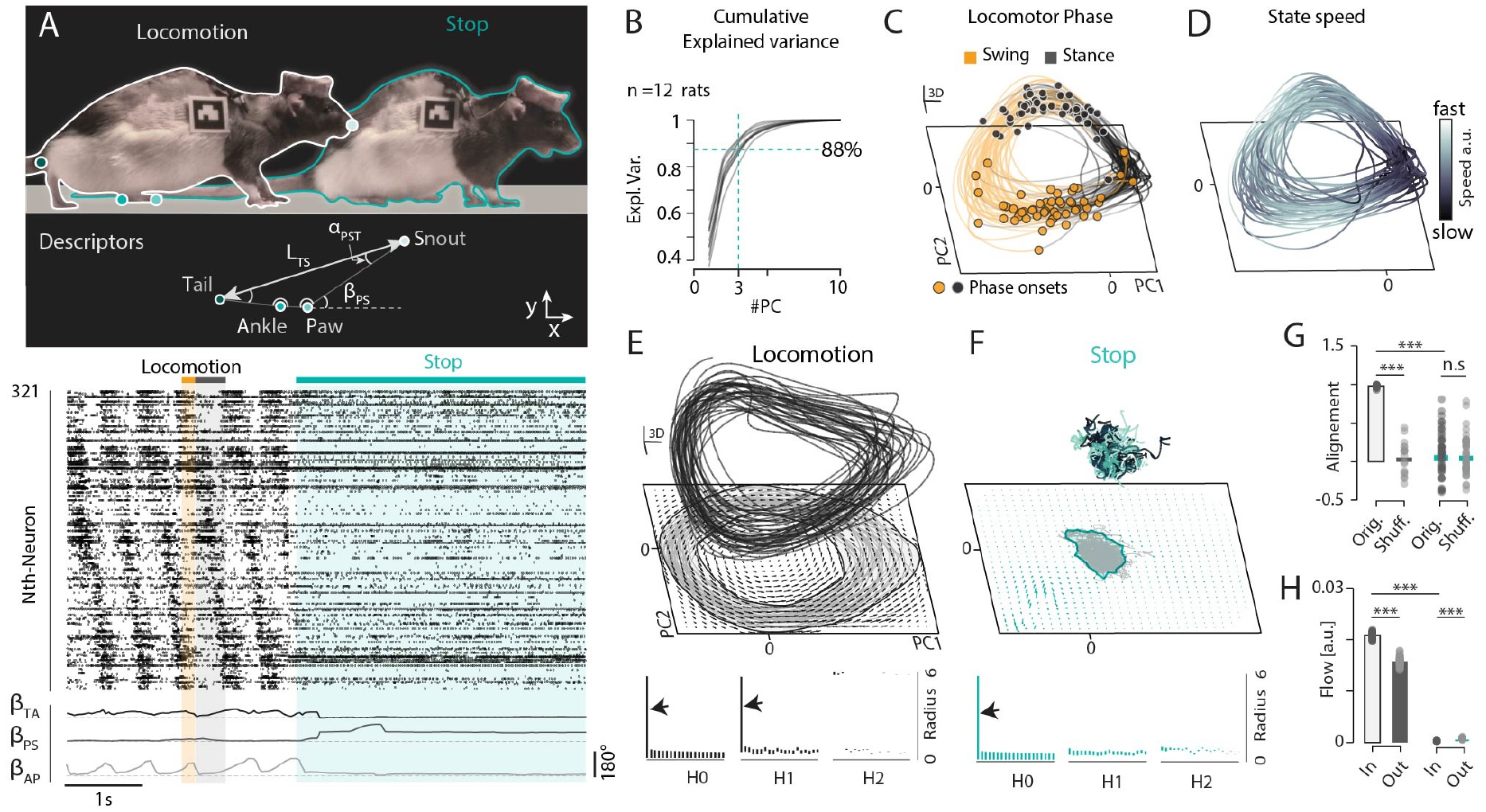
From walk to stop and the associated neural states. **a**, top: sample images of walk-to-stop with pose metrics indicated. Bottom: Spinal raster plot from walk-to-stop (phase sorted, stop: cyan). Swing/stance indicated (orange/grey). Pose metrics, bottom. **b**, cumulative explained variance versus number of principal components (x-axis) of the population firing rates (cycle-averaged) is high (PC1-3 explains >80%, broken lines, n=12 animals), which reveals low-dimensionality. **c**, state space trajectories during walking has a ring-like structure, the ‘locomotor manifold’, swing/stance phases indicated. **d**, The movement on the manifold slows down during the stance (speed indicated by grey-scale). **e**, State space flow-field (arrows at bottom) indicates the direction of motion. Persistent homology estimates the manifold as a solid ring (*H*_0_ and *H*_1_ but not *H*_2_ have lines, arrows, bottom)^28^. **f**, Upon stopping the state switch to a non-trivial fixed point attractor. Repeated stops form a postural manifold, which is topologically solid (only one line in *H*_0_, arrow, bottom). **g**, Alignment of trajectories in the locomotor vs. postural manifold, measured using (normalized projection of flow line on mean flow, n=12 animals). **h**, Mean voxel flow inside the manifolds, indicates limit cycle state during locomotion and fixed point states during stopping. Wilcoxon signed-rank test, ***: p<0.001.

### Attractor manifolds

The onset of the walk-to-stop transition is important for uncovering the transient dynamics of the spinal network. Since stopping is controlled by the animal and it is unknown when it decides to stop, it would be beneficial if we could also control stopping. We do this by optogenetic stimulation of PPN in multiple trials to cover all phases of the walking cycle. Following stimulation, spinal activity changes rapidly from sequential to stable tonic firing (three trials are shown, **Fig. 4a**). At the same time, the animal is stopped in various postures, seen as the hind paw velocity is going to zero (bottom, **Fig. 4b**). The state trajectory in latent space rapidly moves from circular motion toward a stable fixed-point where it remains throughout the stimulation period (**Fig. 4c**). Once stimulation stops, population activity resumes sequential activity and the neural trajectory returns to rotational dynamics within the locomotor manifold (**Supplementary Video 2**). Repeated stimulation in trials reveals a larger manifold representing more diverse postural positions. This larger manifold surrounds the volitional stop manifold (83% overlap, pink vs. cyan, **Fig. 4e**). Next, we ask if various postural stops are represented by the same fixed point with some noise, or whether there are distinct points forming a quasi-continuum. Hence, we inspect the movement inside the postural manifold around the fixed-points associated with a stop. The trajectory during steady-state remains in close vicinity of the fixed-point, i.e. the variance of the state is smaller than the size of the manifold (PC1, black vs. pink **Fig. 4f**). This pattern suggests that the postural manifold is composed of a set of distinct attractive fixed-points representing various postural stoppings. The various physical postures are also found both in natural and PPN-induced stops (**Extended Data Fig. 6**).

**Fig. 4.**
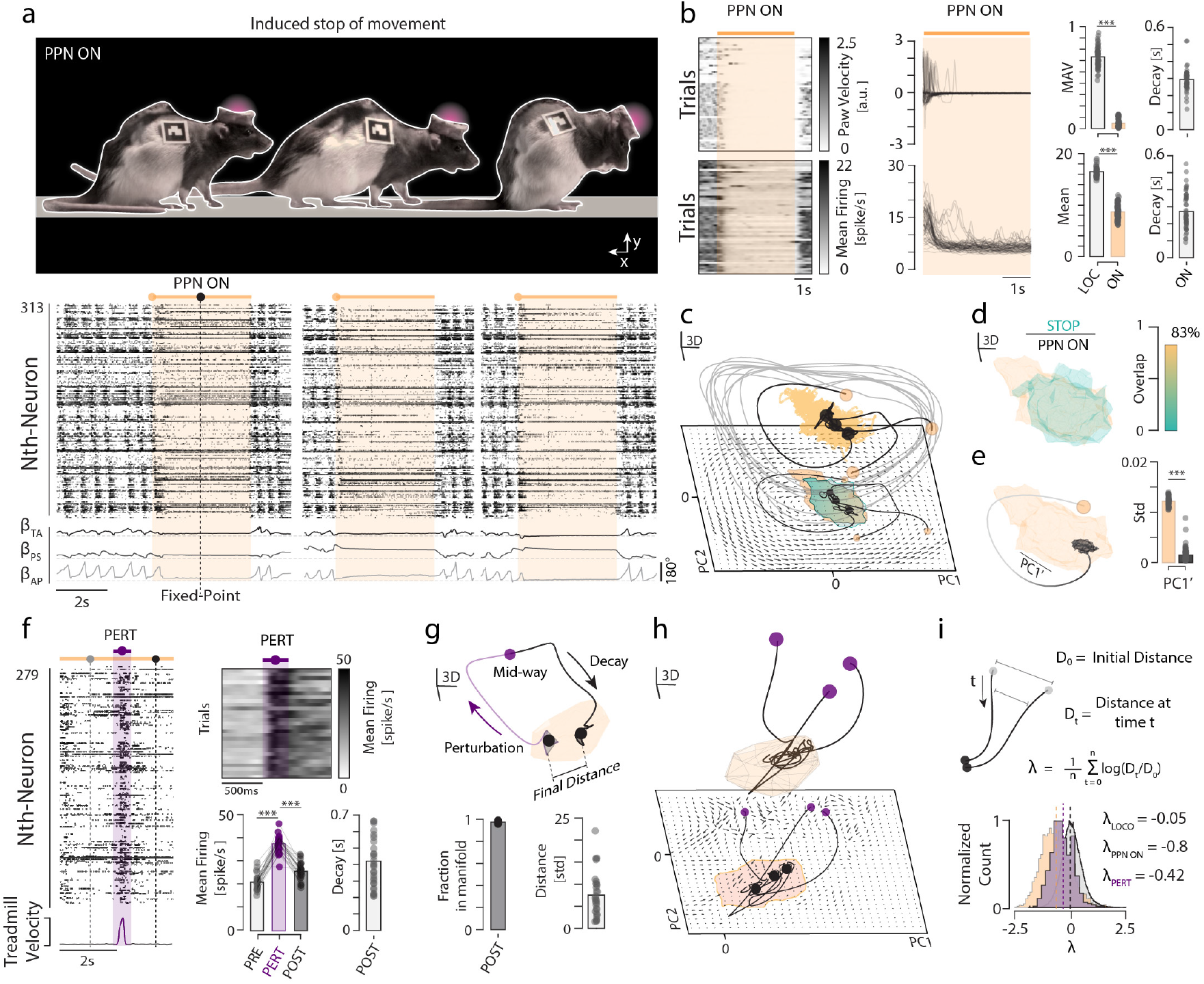
Induced stopping and manifold stability. **a**, Top: Examples of PPN-induced motor arrest: during sitting, walking and grooming. Bottom: Spinal population activity during walk-to-stop transitions (3 trials, PPN-stimulation: yellow shaded, broken line: point of steady-state assumption). Paw kinematics (bottom). **b**, Paw velocity across trials (n=53) and PPN stimulation (“PPN on”, top, left and middle). Mean firing of population across trials during stimulation (bottom, left and middle). Bar graphs: mean absolute paw velocity (MAV), mean firing, and their decay time (“Decay”, rightmost). **c**, In neural state space, walk-to-stop transition show convergence from locomotor manifold (gray trajectory) to postural manifold (yellow region, trials from a, PC1-3). **d**, Postural manifold is larger yet overlaps with that of natural stops (yellow vs. green, 83%). **e**, Postural manifold is larger than random movement around fixed point (yellow vs. black, standard deviation of PC1). **f**, Limb perturbation by jerk of belt velocity during PPN-induced stops (left). Bottom: Belt velocity. Top-right: Mean firing during perturbation (purple) across trials (n=23). Bottom: Bar graphs indicating significant increase/decrease in mean firing and the decay time after perturbation. **g**, Excursion of trajectory associated with perturbation, decays back to a nearby fix point (dots) in manifold (yellow). **h**, three sample perturbations in state space and their trajectories back to the postural manifold (yellow). The flow lines are shown in 2D (bottom). **i**, Attractor property verified by convergence of nearby trajectories (top) using ‘Lyapunov exponents’, *λ*. Histograms of *λ* (bottom) indicate primarily negative values. Wilcoxon signed-rank test, ***: p < 0.001.

A central element of attractor dynamics is stability, which can be verified using perturbations^39^. Now that we can control the cessation, we can also apply perturbations. Hence, half-way through a PPN-induced stop, an impulse of the treadmill belt speed is applied (purple **Fig. 4g** and **Extended Data Fig. 9**). This causes an immediate increase in firing rates, which decays shortly afterwards (∼400 ms, right). This is represented as an excursion in latent space that decays back to a new location within the postural manifold, often several standard deviations from the original position (>5 std, bottom **Fig. 4g**). The postural manifold has attractor properties, which we further verify by comparing trajectories of multiple perturbations (**Fig. 4h**). If two nearby trajectories from different trials converge over time, it is an indication of attractor dynamics (top, **Fig. 4i**). If they diverge, the dynamics is unstable. This is quantified as the exponential decay/growth of the distance, *D*, and the decay/growth rate, *λ*, known as the Lyapunov exponent^40^. A negative/positive *λ* indicates convergence/divergence, and hence the stability of the state. If *λ* is near zero, the states co-evolve and the system is in steady-state, as during walking and the locomotor manifold, which has a near-zero *λ* (*λ*_*loco*_ = −0.05, bottom, **Fig. 4i**). Co-evolving stable periodic orbits are supportive of the theory prediction that the locomotor manifold is composed of a nested set of limit cycles. In conclusion, the walk-to-stop is associated with a transition between two neural attractor states in neural activity, where walking is generated by rotational population dynamics within a ring-shaped manifold and stopping is maintained by a solid postural manifold containing various discrete fixed points. These observations agree with the dynamics of the proposed theory (**Fig. 1**). The exact mechanism of how the transition is accomplished is an open issue, but modulation of cell types that represent aspects of the Mexican hat projectome would certainly fulfill this transition, and experiments on V1 manipulations seem to support it^20,41^.

### Cell types during walk and stop

So far, we have only described the collective dynamics of the spinal network during walking and stopping without regard to cell type. We now consider individual neurons during walking and stopping. During PPN-induced stops, the mean population firing rate decreased, although it did not reach zero (**Fig. 4b**). However, not all cells have a decrease in firing. When sorting neurons according to their steady-state firing rates during PPN-induced stops, we find a continuum of rates from complete silence to an increase (purple vs. black, **Fig. 5a**) across the dorso-ventral axis (**Fig. 5b-d, Extended Data Fig. 10**). To simplify, we define four groups: Neurons that 1) are silent during walking and start firing during stop (‘appear’), 2) fire more during stops than during walk (‘increase’), 3) fire less during stops than walks (‘decrease’), and 4) are silent during stops (‘silenced’, **Fig. 5e-f**). Most (∼70%) decrease their firing or become silent during stopping. The change for all neurons can be visualized by plotting the firing of individual neurons during locomotion (x-axis) versus during stopping (y-axis), where the unity line splits the population into the groups overall (**Fig. 5g**). Next, we ask if PPN-induced stopping has a similar effect on activity compared to volitional stopping. A plot of the change in firing rates for the two conditions reveals that most but not all fall on the identity line green vs. broken lines (**Fig. 5h**). This indicates broadly similar modulation patterns under PPN-induced and volitional stopping, with a subset of neurons showing condition-specific differences.

**Fig. 5.**
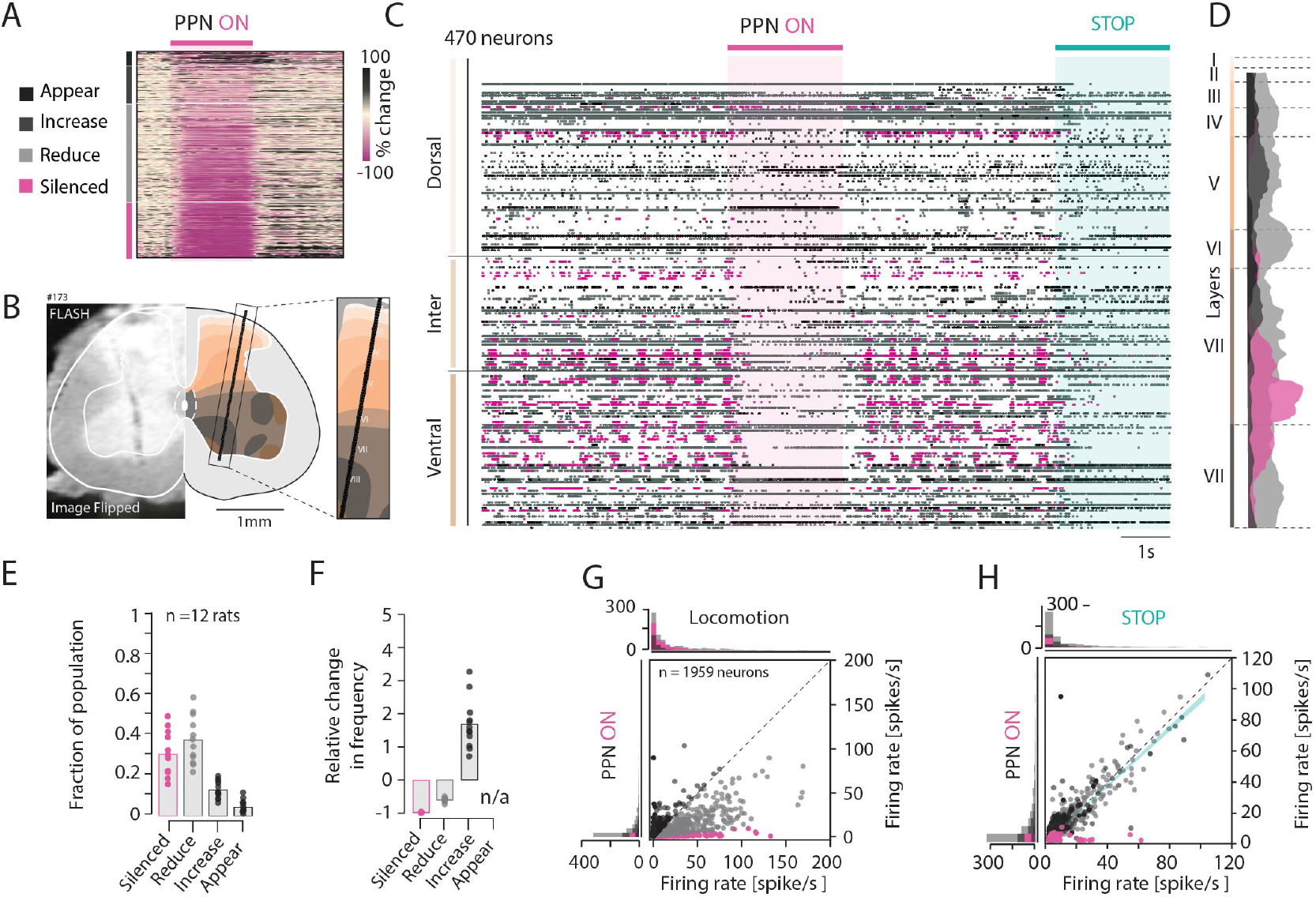
Behavior within neuronal population during walk and stop. **a**, Color coded sample firing rates of neurons before, during and after PPN-induced stops (indicated) ordered according to firing increase during stop (y-axis, PPN-stimulation indicated). Neurons are categorized according to their response: Appear, increase, reduce and silenced during stops. **b**, Neuronal locations across layers are reconstructed from MRI scans (L5 segment). **c**, Sample raster of population spiking with the silenced group highlighted (purple, N=470 neurons). Both PPN-induced and volitional stop indicated (shaded regions, volitional stop: green). **d**, Layer-wise distributions indicate the silenced group is mainly in layers VII-VIII. **e**, Relative fraction of populations within the four neuronal groups, and their relative changes in frequency, **f. g**, Firing rates across population during locomotion (x-axis) versus during induced stop (y-axis) with identity line (broken line) and distributions (top and left). **h**, Absolute change in neuronal firing during PPN-induced (x-axis) and natural arrest (y-axis), with linear fit (green) close to identity line (broken). e-h: N=1959 neurons across n=12 rats.

## Discussion

Neuroscience increasingly uses dynamical systems theory to interpret brain function, especially in large networks of interacting neurons^8,11,42^. This approach is particularly relevant for studying the spinal cord, as movement consists of time-dependent sequential activation of muscles. Here, we address the generation of walking using neural state-space descriptions and dynamical systems theory. It is also well known from dynamical systems theory that a transition between states often gives important insight into systems. For these reasons, we focus not only on the generation of locomotion but also on how the neural system can perform walk-to-stop transitions. First, we develop a theory where motor function arise purely out of the spatial properties of the spinal cord and the primary role of cell types is distinct longitudinal projections, forming local recurrent excitation and longer-range inhibition. Using this concept, we can explain the rotational population activity and how motor rhythms could emerge at the network level without having to rely on cellular pacemaker properties. Traditionally, rhythm generation has been attributed to neuronal pacemakers^15,16,43^. Although such cells have been identified^44^, their necessity for locomotion in mammals remains uncertain^18,45,46^. More critically, it is not clear how these cellular mechanisms can be stopped in various phases by descending control, as seen in experiments^5,47,48^. Hence, investigating the walk-to-stop transition likely provides important mechanistic insight into rhythm generation itself.

Our theory is capable of explaining postural stops. We test these and other model predictions using Neuropixels probes implanted in the lumbar spinal cord of rats that can walk freely, thus also stop at own volition. In addition, we override the generation of spinal motor patterns by stimulating a brain stem nucleus, the PPN, and thus forcing a walk-to-stop transition. Our experiments indicate that during walking the spinal population performs a low-dimensional rotation in neural state space. The trajectory is contained inside a ring-shaped manifold, the ‘locomotor manifold’, which has attractor properties similar to a limit cycle. The walk-to-stop transition occurs by the state changing from a limit cycle dynamics to a stable fixed point inside a separate ‘postural’ manifold. The manifold is composed of many non-trivial fixed points, i.e. they are non-zero, and together they form a quasi-continuum, which is somewhat similar to a continuous attractor network^28,29^. The theory did not support a ring-shaped postural manifold, a prediction that was unexpectedly supported by the experiments (cf. **Fig. 3-4, Supplementary Video 2**). The interpretation of a postural manifold not being ring shaped is that it is not possible to maintain a stable posture in all phases of the walking cycle.

Our observations challenge the conventional assumptions of spinal central pattern generators. First, we observe a continuous sequence of activity, whereas the traditional half-center view has modular activity with reciprocal inhibition that ensures alternation between flexors and extensors^14–16^. Such an alternation scheme predicts activity with only two phases, which seems at odds with the near-uniform phase distribution (**Fig. 1j**). While trying to rescue the traditional scheme by adding more module layers could create additional phases, these layers are connected in a feedforward manner, and thus amounts only to fixed time delays. As a result, the phase differences change as the frequency changes, which is counter-productive to behavior. Constant phase-lags could be gained by sensory feedback^49^, but this is likely not the only mechanism, since experiments on sensory-deprived spinal cords also have wide phase distributions^10,50,51^. A plausible additional explanation is that the architecture of the spinal network is similar to that of many other parts of the nervous system: recurrent and balanced E/I connectivity^52,53^, which ensures robust rotational dynamics through internal negative feedback^10,52,53^.

An interesting experimental observation during postural stops is that interneurons have vastly different activities. We divided the population into four categories: 1) neurons become silent, 2) have reduced firing, 3) have increased firing during postural stops, and 4) cells are inactive during locomotion but become active (appear) during stops (**Fig. 5** and **Extended Data Fig. 8**). These different groups invite speculation of their purpose. Since their activity is no longer needed during a static pose, the group that becomes silent could constitute the core of rhythmogenic neurons that have previously been proposed as central pattern generator cells^15,18^. Their location is in the intermediate to the ventral portion (**Fig. 5d**). However, the majority of cells either reduce their firing or stop firing altogether, which could be explained by activation of the recurrent inhibitory feedback of the model: The bifurcation from the limit cycle dynamics to a fixed point depends on inhibition of the ascending inhibitory neurons. This activity likely serves to maintain stability, posture, or processing sensory feedback.

## Data availability statement

All data presented in this study are available from the corresponding author, RWB, on reasonable request. Please find the relevant code associated with modeling and data analysis in the following github repository (https://github.com/BergLab/ManifoldsofWalking).

## Acknowledgements

This work was supported by The Lundbeck Foundation (no. R366-2021-233, R.W.B.), Novonordisk foundation (no. NNF23OC0082192, R.W.B), EIC Pathfinder Open Project 101130161 - Move2Treat (R.W.B.), and The Swiss National Science Foundation (no. SNF Postdoc.Mobility P500PB_206824, S.K.). The authors thank Dr. Yuki Mori for his assistance in the MRI scanning of the tissue and Palle Koch for technical support.

## Author contributions

J.K., G.H., G.L., K.S., M.C.A.B., R.W.B. performed surgeries. S.K. J.K., K.S., R.W.B. collected the data. S.K. and A.W. performed the model simulations. S.K. A.W. and R.W.B. designed and developed the theory. G.H., R.J.S., M.C.A.B., G.L., S.K., and R.W.B. analyzed the experimental data. R.W.B. conceived the original idea. S.K. and R.W.B. wrote the manuscript.

## Disclosures

The authors declare no conflicts of interests. Funded partially by the European Union. However, views and opinions expressed are those of the author(s) only and do not necessarily reflect those of the European Union or the EIC. Neither the European Union nor the granting authority can be held responsible for them.

## Notes

### Competing Interest Statement

The authors have declared no competing interest.

### Summary of Updates

Minor improved in the text. Supplementary data and methods section now included as separate files.

## References

1. Komi, S. et al. Spatial and network principles behind neural generation of locomotion, DOI: 10.1101/2024.10.03.616472 (2024).

2. Wandler, F. D., Lemberger, B. K., McLean, D. L. & Murray, J. M. Coordinated spinal locomotor network dynamics emerge from cell-type-specific connectivity patterns, DOI: 10.7554/eLife.106658.1 (2025).

3. Jun, J. J. et al. Fully integrated silicon probes for high-density recording of neural activity. Nature 551, 232–236, DOI: 10.1038/nature24636 (2017).

4. Carvalho, M. M. et al. A Brainstem Locomotor Circuit Drives the Activity of Speed Cells in the Medial Entorhinal Cortex. Cell Reports 32, 108123, DOI: 10.1016/j.celrep.2020.108123 (2020).

5. Goñi-Erro, H., Selvan, R., Caggiano, V., Leiras, R. & Kiehn, O. Pedunculopontine Chx10+ neurons control global motor arrest in mice. Nat. Neurosci. 26, 1516–1528, DOI: 10.1038/s41593-023-01396-3 (2023).

6. Kaur, J. et al. Pedunculopontine-stimulation obstructs hippocampal theta rhythm and halts movement. Sci. Reports 15, 17903, DOI: 10.1038/s41598-025-01695-8 (2025).

7. Tello, A. J. et al. Dopamine-sensitive neurons in the mesencephalic locomotor region control locomotion initiation, stop, and turns. Cell Reports 43, 114187, DOI: 10.1016/j.celrep.2024.114187 (2024).

8. Churchland, M. M. et al. Neural population dynamics during reaching. Nature 487, 51–56, DOI: 10.1038/nature11129 (2012).

9. Gallego, J. A., Perich, M. G., Miller, L. E. & Solla, S. A. Neural Manifolds for the Control of Movement. Neuron 94, 978–984, DOI: 10.1016/j.neuron.2017.05.025 (2017).

10. Lindén, H., Petersen, P. C., Vestergaard, M. & Berg, R. W. Movement is governed by rotational neural dynamics in spinal motor networks. Nature 610, 526–531, DOI: 10.1038/s41586-022-05293-w (2022).

11. Perich, M. G., Narain, D. & Gallego, J. A. A neural manifold view of the brain. Nat. Neurosci. DOI: 10.1038/s41593-025-02031-z (2025).

12. Kirk, E. A., Hope, K. T., Sober, S. J. & Sauerbrei, B. A. An output-null signature of inertial load in motor cortex. Nat. Commun. 15, 7309, DOI: 10.1038/s41467-024-51750-7 (2024).

13. Frigon, A. & Gossard, J.-P. Evidence for specialized rhythm-generating mechanisms in the adult mammalian spinal cord. J Neurosci 30, 7061–7071, DOI: 10.1523/JNEUROSCI.0450-10.2010 (2010).

14. McCrea, D. A. & Rybak, I. A. Organization of mammalian locomotor rhythm and pattern generation. Brain Res Rev 57, 134–146, DOI: 10.1016/j.brainresrev.2007.08.006 (2008).

15. Grillner, S. & El Manira, A. Current principles of motor control, with special reference to vertebrate locomotion. Physiol. Rev. 100, 271–320, DOI: 10.1152/physrev.00015.2019 (2020).

16. Kiehn, O. Decoding the organization of spinal circuits that control locomotion. Nat. Rev. Neurosci. 17, 224–238, DOI: 10.1038/nrn.2016.9 (2016).

17. Sengupta, M. & Bagnall, M. W. Spinal Interneurons: Diversity and Connectivity in Motor Control. Annu. Rev. Neurosci. 46, 79–99, DOI: 10.1146/annurev-neuro-083122-025325 (2023).

18. Ren, J. & Gosgnach, S. Localization of Rhythm Generating Components of the Mammalian Locomotor Central Pattern Generator. Neuroscience 513, 28–37, DOI: 10.1016/j.neuroscience.2023.01.013 (2023).

19. Dubuc, R., Cabelguen, J.-M. & Ryczko, D. Locomotor pattern generation and descending control: a historical perspective. J. Neurophysiol. 130, 401–416, DOI: 10.1152/jn.00204.2023 (2023).

20. Gosgnach, S. et al. V1 spinal neurons regulate the speed of vertebrate locomotor outputs. Nature 440, 215–219, DOI: 10.1038/nature04545 (2006).

21. Hayashi, M. et al. Graded Arrays of Spinal and Supraspinal V2a Interneuron Subtypes Underlie Forelimb and Hindlimb Motor Control. Neuron 97, 869–884, DOI: 10.1016/j.neuron.2018.01.023 (2018).

22. Berg, R. W., Chen, M.-T., Huang, H.-C., Hsiao, M.-C. & Cheng, H. A method for unit recording in the lumbar spinal cord during locomotion of the conscious adult rat. J. Neurosci. Methods 182, DOI: 10.1016/j.jneumeth.2009.05.023 (2009).

23. Wu, Y. et al. Ultraflexible electrodes for recording neural activity in the mouse spinal cord during motor behavior. Cell Reports 43, 114199, DOI: 10.1016/j.celrep.2024.114199 (2024).

24. Komi, S. A., Kaur, J. & Berg, R. W. Spatio-temporal population dynamics of the lumbar spinal cord during locomotion in awake rats. In Program No. PSTR155.13. 2023 Neuroscience Meeting Planner. Washington, D.C.: Society for Neuroscience (2023).

25. Bush, N. E. & Ramirez, J.-M. Latent neural population dynamics underlying breathing, opioid-induced respiratory depression and gasping. Nat. Neurosci. 27, 259–271, DOI: 10.1038/s41593-023-01520-3 (2024).

26. Wimalasena, L. N., Pandarinath, C. & AuYong, N. Spinal interneuron population dynamics underlying flexible pattern generation. Nat. Commun. 16, 9634, DOI: 10.1038/s41467-025-64629-y (2025).

27. Zhang, K. Representation of spatial orientation by the intrinsic dynamics of the head-direction cell ensemble: a theory. The J. Neurosci. 16, 2112–2126, DOI: 10.1523/JNEUROSCI.16-06-02112.1996 (1996).

28. Gardner, R. J. et al. Toroidal topology of population activity in grid cells. Nature 602, 123–128, DOI: 10.1038/s41586-021-04268-7 (2022).

29. Kim, S. S., Rouault, H., Druckmann, S. & Jayaraman, V. Ring attractor dynamics in the Drosophila central brain. Science 356, 849–853, DOI: 10.1126/science.aal4835 (2017).

30. Bácskai, T., Rusznák, Z., Paxinos, G. & Watson, C. Musculotopic organization of the motor neurons supplying the mouse hindlimb muscles: a quantitative study using Fluoro-Gold retrograde tracing. Brain Struct. Funct. 219, 303–321, DOI: 10.1007/s00429-012-0501-7 (2014).

31. Avaltroni, P., Cappellini, G., Sylos-Labini, F., Ivanenko, Y. & Lacquaniti, F. Spinal maps of motoneuron activity during human locomotion: neuromechanical considerations. Front. Physiol. 15, DOI: 10.3389/fphys.2024.1389436 (2024).

32. Flynn, J. R. et al. Anatomical and Molecular Properties of Long Descending Propriospinal Neurons in Mice. Front. Neuroanat. 11, DOI: 10.3389/fnana.2017.00005 (2017).

33. Sengupta, M., Bertram, A., Zhu, S. I., Goodhill, G. J. & Bagnall, M. W. V2b Neurons Act via Multiple Targets to Produce in Phase Inhibition during Locomotion. The J. Neurosci. 45, e1530242025, DOI: 10.1523/JNEUROSCI.1530-24.2025 (2025).

34. Bikoff, J. B. et al. Spinal Inhibitory Interneuron Diversity Delineates Variant Motor Microcircuits. Cell 165, 207–219, DOI: 10.1016/j.cell.2016.01.027 (2016).

35. Kathe, C. et al. The neurons that restore walking after paralysis. Nature 611, 540–547, DOI: 10.1038/s41586-022-05385-7 (2022).

36. Steinmetz, N. A. et al. Neuropixels 2.0: A miniaturized high-density probe for stable, long-term brain recordings. Science 372, DOI: 10.1126/science.abf4588 (2021).

37. Li, G., Komi, S., Sorensen, J. F. & Berg, R. W. A Real-Time Vision-Based Adaptive Follow Treadmill for Animal Gait Analysis. Sensors 25, 4289, DOI: 10.3390/s25144289 (2025).

38. Borgognon, S. et al. Regional specialization of movement encoding across the primate sensorimotor cortex. Nat. Commun. 16, 5729, DOI: 10.1038/s41467-025-61172-8 (2025).

39. Khona, M. & Fiete, I. R. Attractor and integrator networks in the brain. Nat. Rev. Neurosci. 23, 744–766, DOI: 10.1038/s41583-022-00642-0 (2022).

40. Engelken, R., Wolf, F. & Abbott, L. F. Lyapunov spectra of chaotic recurrent neural networks. Phys. Rev. Res. 5, 043044, DOI: 10.1103/PhysRevResearch.5.043044 (2023).

41. Falgairolle, M. & O’Donovan, M. J. V1 interneurons regulate the pattern and frequency of locomotor-like activity in the neonatal mouse spinal cord. PLOS Biol. 17, e3000447, DOI: 10.1371/journal.pbio.3000447 (2019).

42. Durstewitz, D., Koppe, G. & Thurm, M. I. Reconstructing computational system dynamics from neural data with recurrent neural networks. Nat. Rev. Neurosci. 24, 693–710, DOI: 10.1038/s41583-023-00740-7 (2023).

43. Ausborn, J., Snyder, A. C., Shevtsova, N. A., Rybak, I. A. & Rubin, J. E. State-dependent rhythmogenesis and frequency control in a half-center locomotor CPG. J. Neurophysiol. 119, 96–117, DOI: 10.1152/jn.00550.2017 (2018).

44. Brocard, F. et al. Activity-Dependent Changes in Extracellular Ca2+ and K+ Reveal Pacemakers in the Spinal Locomotor-Related Network. Neuron 77, 1047–1054, DOI: 10.1016/j.neuron.2013.01.026 (2013).

45. Dougherty, K. J. & Ha, N. T. The rhythm section: an update on spinal interneurons setting the beat for mammalian locomotion. Curr. Opin. Physiol. 8, 84–93, DOI: 10.1016/j.cophys.2019.01.004 (2019).

46. Koronfel, L. M., Kanning, K. C., Alcos, A., Henderson, C. E. & Brownstone, R. M. Elimination of glutamatergic transmission from Hb9 interneurons does not impact treadmill locomotion. Sci. Reports 11, 16008, DOI: 10.1038/s41598-021-95143-y (2021).

47. Bouvier, J. et al. Descending Command Neurons in the Brainstem that Halt Locomotion. Cell 163, 1191–1203, DOI: 10.1016/j.cell.2015.10.074 (2015).

48. Capelli, P., Pivetta, C., Esposito, M. S. & Arber, S. Locomotor speed control circuits in the caudal brainstem. Nature 551, 373–377, DOI: 10.1038/nature24064 (2017).

49. Pazzaglia, A. et al. Balancing central control and sensory feedback produces adaptable and robust locomotor patterns in a spiking, neuromechanical model of the salamander spinal cord. PLOS Comput. Biol. 21, e1012101, DOI: 10.1371/journal.pcbi.1012101 (2025).

50. Tresch, M. C. & Kiehn, O. Coding of locomotor phase in populations of neurons in rostral and caudal segments of the neonatal rat lumbar spinal cord. J. neurophysiology 82, 3563–74 (1999).

51. Kwan, A. C., Dietz, S. B., Zhong, G., Harris-Warrick, R. M. & Webb, W. W. Spatiotemporal dynamics of rhythmic spinal interneurons measured with two-photon calcium imaging and coherence analysis. J Neurophysiol 104, 3323–3333, DOI: 10.1152/jn.00679.2010 (2010).

52. Berg, R. W., Willumsen, A. & Lindén, H. When networks walk a fine line: balance of excitation and inhibition in spinal motor circuits. Curr. Opin. Physiol. 8, 76–83, DOI: 10.1016/j.cophys.2019.01.006 (2019).

53. Lindén, H. & Berg, R. W. Why Firing Rate Distributions Are Important for Understanding Spinal Central Pattern Generators. Front. Hum. Neurosci. 15, 719388, DOI: 10.3389/fnhum.2021.719388 (2021).

