## Supplementary Information for "Neural manifolds that orchestrate walking and stopping"

### Methods

In this Methods section, we describe the experimental protocols, data analysis, and the details of our computational modeling. First, we start with our theory of central pattern generation in the spinal cord associated with the generation of walking and how the spinal network can perform stopping while supporting posture.

#### Theory of spinal network function

Contrary to traditional models of central pattern generation, which are not based on spatial features, we propose that the location of interneurons in the spinal cord and their connectivity patterns form the basis for the generation of locomotion in mammals[1] as well as in aquatic species[2,3]. A central part of this rational is that traditional models are modular and not able to generate rotational population activity, which are solid experimental observations in previous studies[4–7] and in the current investigation (**Fig. 2**). This essential discrepancy between experiment and theory is further problematic in explaining how an animal or a human can stop walking in any part of a cycle. That an animal can stop moving at any phase is obvious when observing their behavior. We propose that rotational population dynamics and the capacity to stop in any phase are two sides of the same coin. Hence, when forming a theory of spinal motor function, it is essential that models possess rotational dynamics[4]. Here, we adopt and adapt the recently developed theoretical framework in which rhythm-generation and rotational dynamics are achieved by networks with recurrent excitation and inhibition[1,4]. In this approach interneuronal connectivity, or "projectome", forms a Mexican hat, i.e. local recurrent excitation and longer range inhibition, in the longitudinal direction of the spinal cord. If the lobes of the projectome are asymmetric, traveling waves arise from the spinal cord during locomotion. This idea is not new and has been proposed and demonstrated in various species[8–12]. The propagation of waves in limbed vertebrates has also been demonstrated, for example in frogs[13], salamanders[14], and newborn rodents[15,16], which could also explain a wave within motoneuron pools[16–18]. Systemic block of inhibition leads to a fast seizure-like wavefront that effectively synchronizes nerve activity[19]. Surface field potentials show sinusoidal waves in cats under fictive rhythmic scratching[20,21]. However, another report was unable to support such a wave among interneuron activity[22], suggesting that the issue could be more complicated and even specific to cell types if it is a propagating wave. This issue remains to be further investigated in future studies. Here, we adapt the structural mechanism described in a previous report[1], to investigate rhythm-generation and the capacity to stop.

**Spinal network as a simplified 1D model** We adapt our model into a simplified one-dimensional (1D) model to mimic the spinal network along the rostro-caudal (RC) axis, while preserving the key features of cellular composition and asymmetric projection biases (**Fig. 1a-b**). Using this 1D framework, we find that selective inhibition of the interneurons that constitute aspects of the projectome could induce rhythm arrest, characterized by a transition from oscillatory activity to tonic firing (**Fig. 1d**). The particular interneurons responsible for these aspects are not known currently, but could consist of V1 and V2b for ascending and descending inhibition, respectively, or V2a cells for local excitation. As we observe (**Fig. 1d-j**) the modulation (silencing or activation) of these cells has similar effects as observed in experiments, hence support our theory.

The details of the implementation of this theory are the following. Spatially distributed networks were constructed by placing $n = 500$ neurons on a line, assigning each a scalar spatial coordinate between 1 and $n$. For simplification, we assume only five subpopulations, although more are likely to be involved in the generation of motor activity. We defined the identity of the subpopulation $p(i)$, with $p \in [1..5]$ and $i \in [1,n]$ , randomly assigned with fractions $f_p$, which satisfies the same excitation and inhibition. Network connectivity was instantiated by assigning each subpopulation a Gaussian synapse probability, $P_{syn}(i,j)$, defined over pairwise distances $d_{ij} \in [-n,n]$. The probability of synapsing of the neuron $j$ with the neuron $i$ was thus defined by

$$P_{syn}(i,j) = G_p \mathcal{N}(d_{ij} | \mu^{p(j)}, \sigma^{p(j)})$$

with $G_p$ being a scaling factor, producing a sparsity $\approx 70\%$. The values chosen for the mean, $\mu$, and standard deviation, $\sigma$, for the five populations are shown (**Extended Data Table 1**). To include an asymmetry in the projections within the network, the means were assumed to include a caudal bias, seen as more positive values for the populations projecting caudally than rostrally (sp5-direction, **Extended Data Table 1**). From these probabilities, a binary adjacency matrix $\mathbf{A}$ was constructed through the binomial sampling process

$$\mathbf{A}(i,j) \sim \text{Binomial}(1, P_{syn}(i,j)).$$

where $\mathbf{A}(i,j) = 1$ defines a synaptic connection from neuron $j$ to $i$. The hadamard product of $\mathbf{A}$ and the synaptic weight matrix, $\mathbf{G}(i,j) \sim \mathcal{N}(1,0.1)$ yielded the connectivity matrix

$$\mathbf{W} = \mathbf{G} \circ \mathbf{A}$$

**Extended Data Table 1.** The mean, $\mu$, and standard deviation, $\sigma$, of the projection probability for the five subpopulation, and their fraction of the network. The asymmetry of the projectome is reflected in higher values of $\mu$ for the caudally projecting subpopulations (sp3-5) than the rostrally projecting (sp1-2).

| | sp1 | sp2 | sp3 | sp4 | sp5 |
|---|---|---|---|---|---|
| Transmitter | Exc. | Inh. | Exc. | Inh. | Exc. |
| $f$ | 0.10 | 0.25 | 0.30 | 0.25 | 0.10 |
| $\mu$ | -335 | -145 | +5 | +155 | +345 |
| $\sigma$ | 5.7 | 7.2 | 8.3 | 7.2 | 5.7 |

where excitatory (inhibitory) columns of $\mathbf{W}$ were made strictly nonnegative (nonpositive). Finally, a detailed excitatory/inhibitory balance was imposed on connectivity as the values of the rows adding up to zero, i.e., $\mathbf{W}_{i,:}^T \mathbf{1}_n = 0$ for all postsynaptic neurons $i$.

The networks were simulated according to the firing rate formalism

$$\tau \dot{\mathbf{x}} = -\mathbf{x} + \mathbf{W}\psi(\mathbf{x}) + \mathbf{I}$$

where $\tau$ is the $n$ dimensional vector of time constants, $\mathbf{x}$ is a vector with the input potentials of all neurons, $\dot{\mathbf{x}}$ its time derivative, and $\mathbf{I}$ an external input composed of deterministic and stochastic terms, $\mathbf{I} = \mathbf{I}_{det}(t) + \varepsilon(t)$. The noise term is Gaussian with $\varepsilon(t) \sim \mathcal{N}(\mu, \sigma)$. $\psi(\mathbf{x})$ is a sigmoidal transfer function (using the hyperbolic tangent function), mapping the input to the firing rate:

$$\psi(x) = \begin{cases} T \cdot \tanh\left(\frac{g(x-T)}{T}\right) + T, & \text{for } x \leq T \\ f_{\max} \cdot \tanh\left(\frac{g(x-T)}{f_{\max}}\right) + T, & \text{for } x > T. \end{cases}$$

The output firing rate is non-negative and $T$ represents the inflection point of the sigmoid where $\frac{d\psi(x)}{dx} = g$. To capture the natural heterogeneity in neuronal properties, we sampled function parameters and external input probabilistically for each neuron. The baseline Gaussian moments are summarized in **Extended Data Table 2**. The networks were simulated according to the Euler rule, $x_{t+dt} = x_t + \dot{x}_t dt$ with time step $dt = 1ms$.

**Extended Data Table 2.** Mean and variance of the single neuron dynamical parameters of the model. $\tau$ is the firing rate time constant. $g$ is the input gain from presynaptic activity. $T$ is the inflection point of the neuronal transfer function. $f_{max}$, together with $T$, gives the asymptotic maximal firing rate. $\varepsilon$ is the additive noise term of the external drive to neurons. $I_{det}$ is the deterministic external drive to neurons.

| | $\tau$ | $g$ | $T$ | $f_{max}$ | $\varepsilon$ | $I_{det}$ |
|---|---|---|---|---|---|---|
| $\mu$ | 8 | 1 | 40 | 40 | 0 | 40 |
| $\sigma$ | 0 | 0.05 | 2 | 2 | 0.5 | 2 |

**Model Dynamics Analysis** The model networks were simulated to investigate their transition from rhythmic to stationary activity. The networks were simulated for $375s$ total, split into 25 segments of $15s$ each. Each segment consisted of a $12s$ rhythmic "locomotor period" followed by a $3s$ stationary "arrest period". For this family of networks, increasing the gain of $sp_2$ or decreasing that of $sp_4$ tends to increase the oscillatory frequency[1]. Under the hypothesis that a *set* of unique limit cycles could produce a set of unique fixed points upon arrest induction, we allowed the mean gain, $\mu(g)$, to vary between the trial segments. For each trial, $\mu(g)$ for the subpopulations $sp_2$ and $sp_4$ were exponentially sampled from $[0.9, 1.3]$ producing a rhythmic state with a unique cycle period within the behavioral range. To induce arrest, the mean external input, $\mu(I_{det})$, was varied to each subpopulation until the network rhythms were reliably terminated. For $sp_2$, $sp_3$ and $sp_4$, rhythm arrest was induced by changing the mean external input of 40 by $-18$, $-12$ and $+21$, respectively.

To more accurately compare model dynamics with experimental data (**Fig. 1d**), simulated firing rates were normalized and converted to spike trains through the inhomogeneous Poisson process

$$S_n \sim \text{Poisson}(r_n \Delta t),$$

with $S_n$ being the spike count in the time bin $n$, $r_n$ the min-max normalized firing rate, and $\Delta t$ the bin size (here $\Delta t = 1ms$). The spikes were sampled independently across the bins using a Poisson distribution random number generator (`poissrnd.m` in MATLAB). Binary spikes were converted back into activity traces by convolution with the Gaussian kernel, (similar to the analysis of experimental data - see below)

$$k(t) = \frac{1000}{\sqrt{2\pi}\,\theta} \exp\left(-\frac{t^2}{2\theta^2}\right),$$

defined in the interval $t \in [-4\theta, 4\theta]$ with $\theta = 25$. The convolution $\tilde{r}_i(t) = (k * S_i)(t)$, with $\tilde{r}_i(t)$ being the filtered activity of neuron $i$ at time $t$ and $S_i$ its spike train, was performed using the `conv.m` function in MATLAB. To build a principal component space for further analysis and visualization, $\tilde{r}_i(t)$ was normalized over time and all cycles were extracted from locomotor periods within a simulation. Cycles were extracted by first performing principal component analysis on the normalized activity traces and then smoothing the scores along the leading principal component using functions `pca.m` and `movmean.m` with a 20ms window in MATLAB. Peaks in the signal along PC1 were detected with `findpeaks.m`, using a minimum peak height- and prominence of 2 standard deviations and minimum peak distance of 200 ms. Peak distances with a longer period than 1000 ms, or falling within the arrest period, were discarded. The remaining candidates were indexed and stored as locomotor cycles. From the extracted cycles, a single mean cycle was computed by averaging. From this mean cycle, the three leading principal components were computed again with `pca.m`. This 3D space was used as a projection space for all further analysis (**Fig. 1g-j**). For both locomotor and arrest periods, persistent barcodes for homology groups $H_0$ , $H_1$ and $H_2$ were computed from the latent traces using an open source persistent homology package `Ripser`[23] for Python.

#### Manifolds, attractors, and the walk-to-stop bifurcation

A neural 'manifold' is defined as a low-dimensional geometric representation of high-dimensional population activity that - due to some constraints or internal relationship - are confined to a subspace[24,25]. In our investigation, we demonstrate that during walking the spinal population activity is represented by a neural manifold, which has a ring-like structure, the locomotor manifold (**Supplementary Video 1**). Since the ring is solid it is not a torus, contrary to circular manifolds observed in other parts of the nervous system[26]. During the walk-to-stop transition, the state changes from rotational dynamics to a different solid manifold, the postural manifold, which is not ring shaped. Both the postural and the locomotor manifolds work as attractors, since trajectories converge towards them after a walk-to-stop bifurcation or the system is perturbed, as demonstrated by the negative Lyapunov exponents (**Fig. 4i**). Since the locomotor manifold has rhythmic rotational dynamics, we consider it a type of attractor called a limit cycle attractor. A limit cycle attractor has circular motion in a closed-loop trajectory that attracts nearby trajectories over time. These trajectories move asymptotically towards the limit cycle and therefore the limit cycle has no width: it is not a ring, but a circular line. However, if the parameters of the neural system change slightly, for instance, the descending drive or the neuronal gain, the limit cycle will also change slightly. Limit cycles that have non-infinitesimal width are often seen in stochastic biological processes[27]. Hence, the manifold does not strictly have to be a circle to have limit cycle behavior. Since the locomotor manifold is an attractor with periodic activity but a circular shape with width, it could represent a set of nested limit cycles arising from slightly changing parameters over time. The changing parameters could represent several different biological functions. For example, the radius of rotation has been indicated to represent the force of muscles, which is changed from step to step depending on external conditions, and this can be controlled in the model by a constant descending drive[4]. In the current model, the width of the locomotor manifold is partially due to a variance in the neuronal gain among other things (**Extended Data Table 2**). It is important to note that although some of the width of the locomotor manifold is due to noise, the variability of the trajectory inside the manifold is a means of adjusting the step, which in the model is accomplished by neuronal gain modulation, and should therefore be considered a crucial element of motor control. In conclusion, we propose that during walking, the activity of the spinal population is confined to a locomotor manifold, which acts as a limit cycle attractor.

What type of attractor is the postural manifold? Experimentally, we observe the state transition from rotation to a stable fixed point within a large region of the neural state space, which we define as the postural manifold. Each behavioral stop has its own fixed point that is uniquely located in the state space, which represents the particular posture of the animal. The manifold is delimited by repeating the walk-to-stop transition many times so that the set of fixed points forms a quasi-continuum. Since animals and humans can stop in almost any phase of the walking cycle, we expected the postural manifold to cover a large region, perhaps even a ring in state space, to represent posture of all phases. This region works as an attractor and could have continuous attractor properties, as seen in circular systems in other parts of the nervous system, such as the substrate for head-direction and grid-cell representation[26,28]. A continuous attractor network is often imagined as a network of neurons connected in a ring with local recurrent excitation and more global inhibition[29]. A bump of activity can be kept stable at a given location, but moved due to external drive or some temporary asymmetry in the recurrent connections. The movement of a bump could represent the sequential contraction of motoneurons, if connected properly, and a static bump could represent a steady posture during a stop. This is the reasoning for the 1D model of the spinal cord network (**Fig. 1**). In reality, a continuous attractors network is difficult to verify experimentally and may also not be the most stable solution in the motor system. During static posture, there is a constant pull from gravity that would slowly move the bump of activity, and thus the limbs out of their position. Instead, a discrete set of stable fixed points would resist such a pull, which is a better solution. It is also easier to envision a neuronal network with stable fixed points that does not require perfect connection symmetry. In our model of a linear network, there is only one stable fixed point for one set of network parameters. When changing these parameters, the fixed point will have a different location in the state space. This is seen as a collection of fixed points (**Fig. 1h-j**), which we proposed as the postural manifold that we see in experiments (**Fig. 4c**, **Supplementary Video 2**).

#### Experimental methods

##### Animals and Ethical statement

Adult Wild-type Long Evans rats of both sexes (aged 12-24 weeks, 450-600 g) were used to perform the experiments. The rats were obtained from Charles River Laboratories and housed in cages equipped with bedding, food, water, temperature, and air sensors and kept in a 12 h light/ 12 h dark cycle. Before any surgical procedure, the rats were housed in pairs for one week in the animal housing facility for acclimatization. All experimental procedures were carried out in accordance with the ARRIVE guidelines and the Council of the European Union (86/609/EEC). The study was approved by the Danish veterinary and food administration (animal research permission numbers 2019-15-0201-00018 and 2024-15-0201-01739).

##### Surgery

The surgical procedures were performed under aseptic conditions. The rats were anesthetized using gas anesthesia (isoflurane in a mixture of air and oxygen). The surgery room was divided into two sections: a dirty and a clean section. In the dirty section, an initial part of the procedure was performed, such as shaving hair from the surgical area of the rat and cleaning the shaved area with antiseptic and disinfectant liquid (chlorhexidine) in movements from inside to outside, followed by cleaning with ethanol (70%). The rat was moved to the clean section, which is the surgical table with a stereotaxic robot, on a heating pad with a temperature sensor, and a pulse-oximeter (PhysioSuite, Kent Scientific) was used to monitor the state of the animal during surgery. The head of the rat was fixed using the ear bars. An ointment (Ocryl gel) was applied to keep the eyes moist and the surgical area was again rinsed with 70% ethanol. Then a sterile drape was placed on the anesthetized rat, a hole was made on the drape at the surgical area of interest, and a film with a fine cut in the middle (Mepitel, Molnlycke Healthcare, Goteborg, Sweden) was placed on the shaved sterile area of the rat. At this point, the surgery can continue as a brain surgery or as a spinal surgery (see below).

##### Brain Surgery and optogenetics

The skin on the head was cut open and the skin was stretched to increase the visibility of the skull. Bulldog forceps were used to keep the area open. The connective tissue was removed and the skull was scraped with a curved scalpel to remove the periosteum and then dried with electrocautery and hydrogen peroxide (2%). The bone surface was painted with a thin layer of dental cement (C&B Metabond, Parkell). A 3D-printed head plate was glued to the skull with an attached copper mesh[30]. Craniotomy was performed and an AAV virus (AAV9-CamKIIa-ChrimsonR-mScarlet-Kv2.1, which was a gift from Christopher Harvey through Addgene.org[31]) was injected with a glass capillary using a stereotaxic robot (Neurostar GmbH, Tubingen, Germany) into the Pedunculopontine nucleus (500 nl injection volume, coordinates DV: 7.8, ML: 1.58, AP: -7.8). An optical fiber (ø200$\mu$m, 10 mm in length, CFML12L10, Thorlabs, Sweden, or RWD, China) was subsequently implanted with the tip 200 $\mu$m above the injection site and the hole was sealed using cyanoacrylate glue followed by dental cement. The success of implantation was verified both by assessing the behavioral response (stopping movement when optogenetic stimulation) and in a subset of animals (4 out of 12) by subsequent inspection of the optical fiber placement in post mortem magnetic resonance imaging (MRI) scans (**Extended Data Fig. 2**). A crown was built out of the copper mesh using a thin layer of dental cement to protect the protruding optical fiber ferrule[30]. Post-operative care was provided as pain relief via buprenorphine (mixed with Nutella[32], sublingual tablets 0.2 mg crushed powdered and mixed with 1 g of Nutella, dose = 0.4 mg/kg, every 12 h for 3 days) and carprofen (subcutaneously (s.c.), 5 mg/kg, once a day for 5 days), which also worked as an anti-inflammatory. The rat was monitored twice a day for 3 consecutive days, followed by once a day for 7 days. Optogenetic stimulation was performed using a red LED laser (625 nm, M625F2, Thorlabs, Sweden) attached to a patch cable and a connecting sleve on the animal implant. The stimulation was constant light or pulses of 100 Hz with a duty cycle of 50%. The power was adjusted to elicit an appropriate response, but with a maximum output of 13 mW.

##### Spinal surgery, laminectomy and implantation

One month after the expression of the virus transgene, rats were tested for motor arrest behavior with optogenetic stimulation in PPN. Once the behavior was confirmed, a second surgery consisted of laminectomy and the implantation of an electrophysiological recording probe. The probe was the Neuropixels 1.0 probe (Imec, Belgium), which is a CMOS-based silicon probe with a high density of recording sites[33]. Spinal surgery was based on a previously published method[34] with modifications. First, the animal was placed in a stereotaxic frame with isoflurane gas anesthesia mixed with oxygen as described above. The fur on the lower back was shaved off and cleaned with chlorhexidine and ethanol swabs. The skin was opened in the thoraco-lumbar region and the tissue was removed from the dorsum of the vertebrae. The vertebrae were then fixed using spinal clamps mounted on a second stereotaxic frame in reverse position. After fixing the vertebrae, the bone was further cleared of the periosteum using a scalpel and dried using a low-temperature cauterizer. The bone surface was carefully edged using hydrogen peroxide applied with a cotton tip applicator. Then glue (Relyx Unicem 2, 3M) was applied to the surface of the neighboring vertebrae to where the laminectomy was to take place. The laminectomy was performed on the T13 vertebrae (over the spinal segments L3-L5) and the dura was carefully opened and partially removed to get access to the surface of the spinal cord. The Neuropixels probe was glued to a removable base plate that was bolted to a 3D-printed protective house. The house and probe were attached to a rod that was attached to the stereotaxic arm. The arm was angled 30-40$^o$ and the probe was inserted $\sim$3 mm into the spinal cord, with an effective dorsoventral depth of $\sim$2 mm. The exposed part of the spinal cord implantation site was covered with a silicon gel. The house and the area around the implant were supported with dental cement (UniFast Trad, GC America, and Tetric EvoFlow, Ivoclar). The incised skin around the surgical area was glued with cyanoacrylate or stitched with sutures. Post-operative care was provided in a similar way as explained previously in the section. The data acquisition and behavioral experiment was conducted the following day, if the animal was assessed to be able to do it and without any signs of pain. An animal that had demonstrated a positive optogenetic motor arrest response (Rat 173) had also been used in a previous study[35]. An overview of the animals used in the study and their features can be found below (**Extended Data Table 3**).

**Extended Data Table 3.** Overview of animals used in this study for the successful outcome and their parameters. **"Pert."**: Number of instances of stops combined with jerking (perturbations) of the the animal limb position. **"Stops"**: Number of volitional stops analyzed per animal. **"PPN stops"**: Number of stops induced by optogenetic stimulation in the brainstem nucleus (PPN) per animal. **"Units"**: Number of spike-sorted single units in the sample. **"Drift"**: Drift after motion correction (residual drift) in $\mu m$. **"No. Cycles"**: The number of stable locomotion cycles used to construct the embedding space and compute the locomotor manifold. **"MRI res."**: The resolution of the MRI scan ot the spinal cord, as in the voxel size in $\mu m$.

| Animal | Name | Pert. | Stops | PPN Stops | Units | Drift | Probe location | No. cycles | MRI res. ($\mu m$) |
|---|---|---|---|---|---|---|---|---|---|
| 1 | 148 | | 9 | | 225 | | L3 | 4 | 50 |
| 2 | 161 | | 124 | 16 | 213 | 1.8 | L3 | 7 | 50 |
| 3 | 173 | | 114 | 98 | 469 | 1.6 | L4 | 54 | 50 |
| 4 | 174 | | 98 | 75 | 322 | 1.5 | L3 | 10 | 50 |
| 5 | 177 | | 99 | | 120 | 1.3 | | 23 | |
| 6 | 178 | | 29 | | 69 | 1.2 | L3 | 7 | 50 |
| 7 | 182 | | 23 | 37 | 80 | 2 | L5 | 15 | 50 |
| 8 | 183 | | 49 | 25 | 87 | 1.4 | L2 | 12 | 50 |
| 9 | 188 | | 119 | | 93 | 3.8 | L1 | 25 | 50 |
| 10 | 190 | 23 | 4 | 40 | 31 | 2.4 | L4-L5 | 8 | 30 |
| 11 | 192 | 25 | 23 | 45 | 279 | 1 | L5 | 14 | 30 |
| 12 | 193 | 29 | 70 | 92 | 106 | 1.5 | S1 | 8 | 30 |

#### Behavioral experiments

The purpose of the experiments was to record the activity of neuronal populations in the lumbar spinal cord during natural walking and the transition to stop in awake, freely moving rats. The animals moved on a custom adaptive speed treadmill while high density Neuropixels probes recorded lumbar spinal activity. To rapidly interrupt locomotion, a red light opsin was expressed in the pedunculopontine nucleus (PPN) and optogenetic stimulation was used to induce motor arrest in different phases of the step cycle.

**Closed-loop experimental system and timing** The experimental setup (**Extended Data Fig. 4**) separated real time treadmill control from high bandwidth acquisition. One computer ("Computer 1") managed treadmill speed. A second computer ("Computer 2") handled data acquisition from the Neuropixels acquisition device (NiDaq) at 30 KHz sampling rate and all digital/analog signals collected at a 20 KHz sampling rate using another acquisition system (RHD 2000 USB interface board, Intan Tech inc.). The Intan acquisition system provided a common hardware clock and a digitized treadmill, triggers, and stimulation signals.

While the rat walked on the treadmill the belt speed was continuously adjusted to match the animal's preferred walking speed. The first camera ("Cam 1", **Extended Data Fig. 4**) mounted in front of the treadmill streamed images to the controller computer ("Computer 1"). A synthetic square fiducial marker (ArUco marker) was attached to the flank of the animal. The marker was detected in each frame and its horizontal position in the image was used as a feedback variable for a Proportional-Integral-Derivative controller that updated the belt speed to keep the animal near the center of the field of view. This real time loop allowed the rat to walk voluntarily while maintaining a relatively stable position in the camera frame. The treadmill belt velocity signal was recorded continuously on an analog channel of the Intan board. This provided a synchronized measure of locomotor speed that could be related to neural activity and kinematics. The second camera ("Cam 2") captured full body kinematics from a fixed viewpoint and was used only for offline analyzes. It was operated in hardware triggered mode. A custom trigger unit generated a continuous TTL pulse train at 30.3 Hz. Each pulse simultaneously triggered the behavioral camera to acquire one frame and was recorded as a digital input on the Intan board. The same trigger unit also delivered a single TTL rising edge to start Neuropixels acquisition. This start pulse was recorded by the Intan board, registering the Neuropixels and Intan time bases. Spiking activity from Neuropixels could therefore be shifted into the Intan clock domain and aligned with camera frames, treadmill signals, and stimulation pulses. Lastly, a second programmable trigger unit generated TTL pulses for optogenetic stimulation. Each pulse was sent in parallel to the LED driver and to an Intan digital input, so that laser onset and offset were time stamped on the Intan clock. These markers were used to define epochs for subsequent analyzes. The synchronization between camera and Neuropixels recording was verified offline, by extracting the light pulses recorded in the video and comparing those events with the trigger events sent to the LED. The uncertainty in the video frame capture and the recorded signals, due to occasional frame skippage or duplication related to, e.g., buffering limitations, was estimated at approximately SEM =2.6 ms, which is a mere 2% of a rat walking cycle event of ~500 ms. Therefore, this uncertainty did not influence the conclusions of the study (see the section "Temporal alignment and uncertainty" below).

**Induced stopping and treadmill perturbations** During behavioral sessions, optogenetic PPN stimulation (40Hz, 50% duty cycle) was administered while the rats walked on the speed-adaptive treadmill. The intensity of stimulation was adjusted for each animal to induce a strong and reliable behavioral stop. Stimulation TTLs recorded on the Intan board defined the onset and offset of optogenetically induced stops. To investigate the stability of the postural state, brief "jerks" of the treadmill belt were applied during the steady state portion of PPN-induced stops. A jerk was given once during a stop and consisted of a 100-ms square monophasic velocity pulse of the treadmill belt. These jerks were used to define perturbation and recovery epochs for neural and kinematic analyzes (**Fig. 4f-i**, **Extended Data Fig. 9**).

##### Magnetic Resonance Imaging

Once behavioral experiments were completed, the brain and spinal cord were preserved for the purpose of post-mortem anatomical reconstruction and verification of the location of the electrode and optical fiber location (**Figs. 2b** and **Ext. data Figs. 2-3**) as described in a previous report[35]. This was performed using structural magnetic resonance imaging (MRI) in the following steps. After transcardial perfusion with phosphate-buffered saline (PBS) followed by paraformaldehyde (PFA, 4%), the tissue was kept overnight in PFA followed by transfer to sucrose solution (30% w/v) and finally to PBS for storage. Upon scanning, the brain and spinal cord were transferred to a Falcon tube (50 ml) containing fluorinert (FC-40, Sigma-Aldrich) to reduce the background signal from the surrounding liquid. The scanner was a 9.4 Tesla pre-clinical magnetic resonance imaging scanner (BioSpec 94/30; Bruker Biospin, Ettlingen, Germany) equipped with a 1.5 T/m gradient coil.

High spatial resolution three-dimensional (3D) structural images were acquired using a constructive interference in steady state (CISS) sequence. The CISS acquisition consisted of four 3D TrueFISP volumes (True Fast Imaging with Steady-State Precession) obtained with orthogonal phase-encoding directions (repetition time (TR) = 7 ms, echo time (TE) = 3.5 ms, flip angle (FA) = $30^o$, and isotropic spatial resolution of 50 $\mu$m). In addition, to facilitate visualization of electrode tracks, T2*-weighted images were additionally acquired using a 3D fast imaging sequence with steady-state precession (3D-FISP) (TR/TE = 7/3.5 ms, FA = $7^o$, and isotropic spatial resolution of 50 $\mu$m). Each 3D TrueFISP volume was motion-corrected using Advanced Normalization Tools (ANTs[36]). The final CISS image was generated by computing an averaged maximum-intensity projection from the motion-corrected TrueFISP volumes, effectively reducing banding artifacts. The resulting CISS volume was then co-registered to the 3D-FISP volume. The resolution was isotropic and was 30 or 50 $\mu$m depending on the length of the scans, 60 or 12 hours, respectively.

#### Data analysis

All analyzes were performed in MATLAB (R2023a, MathWorks) and Python using custom and open-source code.

**Common temporal basis and resampling** To align the different data flows, all data modalities (not videos) were upsampled to the Neuropixels time grid with a sampling frequency $f_{\text{Neuropixels}} = 30$kHz to avoid losing information in the Neuropixels spike times. The result was a shared discrete time axis on which spikes, firing rates, treadmill velocity, kinematics, and stimulation markers were defined, allowing for consistent alignment. After all preprocessing steps (spike sorting, tracking, kinematics, firing rate reconstruction), all data were downsampled to a 1 kHz time base for further analyzes.

**Spike sorting and unit selection** The Neuropixels-recorded data were spike sorted using open source software (Kilosort4[37]), followed by manual curation. Drift correction was applied during sorting to compensate for probe movements relative to tissue. Units were retained as single units if they had a clear refractory period (fraction of inter spike intervals < 1 ms < 0.5%), stable waveform amplitude over the session, and good isolation by visual inspection using default Kilosort4 parameters with drift correction enabled (**Extended Data Fig. 5**).

**Pose estimation and center of mass (CoM) relative kinematics** Behavioral-camera videos were processed with markerless pose-estimation software (DeepLabCut[38], `https://www.mackenziemathislab.org/deeplabcut`) to track the snout, tail base, hind ankle, and paw on the side ipsilateral to the spinal implant (**Extended Data Fig. 6**). All resulting kinematics were resampled and interpolated using an interpolant method that preserves shape by creating a smooth, continuous, cubic spline curve through the data points without introducing oscillations ("pchip.m") at 1 kHz, then filtered using a third-order Butterworth bandpass filter in the [0.1, 10] Hz frequency band. To describe limb motion relative to the body rather than the treadmill, all tracked positions were expressed in a relative coordinate system of center mass (CoM). Let s(t) and b(t) be the 2D positions of the snout and tail base at frame time t. The CoM position was defined as

$$c(t) = \frac{s(t) + b(t)}{2}$$

For a tracked point $p(t)$, the CoM-centered coordinate was

$$\tilde{p}(t) = p(t) - c(t)$$

**Kinematic angles** In addition to CoM-relative positions, we extracted two types of planar angles from the tracked landmarks (**Extended Data Fig. 6**). First, we define segment–segment angles $\alpha_{p_1 p_2 p_3}$ directly from three tracked points. For any triplet $(p_1, p_2, p_3)$, we computed the segment vectors $\mathbf{v}_1 = p_1 - p_2$ and $\mathbf{v}_2 = p_3 - p_2$ in the CoM-centered frame and defined

$$\alpha_{p_1 p_2 p_3} = \arccos\left(\frac{\mathbf{v}_1 \cdot \mathbf{v}_2}{\|\mathbf{v}_1\| \, \|\mathbf{v}_2\|}\right)$$

which gives the angle at point $p_2$ between the two segments. In our notation, we use only the first letter of each landmark: $s$ (snout), $t$ (base of the tail), $a$ (ankle), $p$ (paw). Thus, for example, $\alpha_{sap}$ denotes the angle in the ankle between the segments from the snout to the ankle and from the paw to the ankle, and $\alpha_{tap}$ is the corresponding angle built from the base of the tail, the ankle, and the paw. Second, we calculate segment orientation angles $\beta_{p_1 p_2}$, which describe the orientation of a single segment relative to the horizontal (rostro–caudal) axis. For each segment vector, $\mathbf{v} = p_2 - p_1$ we define

$$\beta_{p_1 p_2} = \text{atan2}\,(w_y, w_x)$$

the signed angle of the segment with respect to the horizontal unit vector. In our compact notation, $\beta_{ts}$ gives the orientation of the body axis (tail-base to snout), and $\beta_{ap}$ gives the orientation of the distal hindlimb segment (ankle to paw) relative to the treadmill axis. Both angle types were computed for each frame and, where relevant, mapped to the gait cycle phase using the same cycle boundaries as for neural phase analyzes.

**Gait event detection and Gait cycle selection** The rostro-caudal component of the CoM-centered paw position, $\bar{x}_{\text{paw}}(t)$, obtained by projecting $\tilde{p}_{\text{paw}}(t)$ onto the longitudinal axis of the treadmill, was used to detect gait events and quantify step cycles. Foot-off (swing onset) was defined as a local minima corresponding to the maximal caudal extension of the paw. Foot-strike (stance onset) was defined as local maxima, corresponding to maximum cranial extension. Gait cycle $k$ was defined as the interval $[t_k, t_{k+1})$ between successive foot-off events. Cycles of irregular duration ($<$ 100ms or $>$ 750ms) or missing events (no foot strike following a foot off). The remaining “good cycles” were used for subsequent analyzes.

**Gait Cycle Normalisation** To compare cycles of different durations, each gait cycle $k$ with foot-off times $t_k$ and $t_{k+1}$ was mapped to a normalized phase

$$\varphi(t) = 2\pi \frac{t - t_k}{t_{k+1} - t_k}, \quad t_k \le t < t_{k+1},$$

The kinematic trajectories were resampled onto a fixed grid of phase bins (e.g., 2000 bins over $[0, 2\pi)$). This yielded phase-normalized kinematic profiles that could be averaged across cycles.

**Firing-rate estimation from spike trains** All analyzes used continuous firing-rate estimates obtained by convolving spike times with a Gaussian kernel. This provides a smooth estimate of the firing probability that varies over time while preserving temporal structure. For neuron $i$, the spike train is

$$S_i(t) = \sum_n \delta(t - t_{i,n}),$$

where $t_{i,n}$ are spike times. The firing rate $r_i(t)$ is defined as the convolution of $S_i(t)$ with a Gaussian kernel $k(t)$:

$$r_i(t) = (k * S_i)(t) = \int k(\tau) S_i(t - \tau)\, d\tau = \sum_n k(t - t_{i,n}),$$

with

$$k(t) = \frac{1}{\sqrt{2\pi}\sigma} \exp\left(-\frac{t^2}{2\sigma^2}\right),$$

where $\sigma = 25$ms is the kernel width that controls the temporal smoothing. $r_i[t]$ is obtained by evaluating this sum at each time point and truncating the kernel to a finite window around each spike. These rates were used consistently across all analyzes for locomotion, natural stopping, PPN-induced stopping, and perturbation epochs. Before dimensionality reduction, firing rates were soft-normalized across time for each neuron to minimize the contribution of neurons showing relatively low firing rates.

##### Phase estimation and step-cycle representation

Given foot-off times $t_k$ and $t_{k+1}$, any neuronal spike $i$ that occurred at time $t$ in the cycle $k$ was assigned a phase

$$\phi(t) = 2\pi \frac{t - t_k}{t_{k+1} - t_k}.$$

For each neuron, the distribution of $\phi$ values between cycles was used to compute the preferred phase (circular mean). These phase values were also used to construct phase-sorted rasters and cycle-averaged firing curves.

##### Temporal alignment and uncertainty

To quantify the timing uncertainty on kinematics introduced by video sampling and handling, we employed the PPN stimulation as a temporal anchor between the video stream and high-bandwidth recordings. For each stimulation trial, we measured

- $t_{\text{TTL}}$: time of the stimulation TTL rising edge on the Intan board.
- $t_{\text{cam}}$: time of the first behavioral-camera frame showing laser light.

The difference

$$\Delta t = t_{\text{cam}} - t_{\text{TTL}}$$

captures the combined effect of frame quantization due to the finite camera frame rate ($f_{\text{cam}} \approx 30.3$ Hz; $\Delta T \approx 33$ ms) and stochastic jitter from camera readout, buffering, OS scheduling, and encoding, which occasionally yielded skipped or duplicated frames. If an event occurs in a continuous $t^*$, it is assigned to the nearest frame time, producing a quantization error $\varepsilon_q \in [-\Delta T/2, \Delta T/2]$. We model each camera-derived event time as

$$\hat{T} = T + \varepsilon_t, \qquad \mathbb{E}[\varepsilon_t] = 0, \quad \text{Var}(\varepsilon_t) = \sigma_t^2,$$

The additional jitter $\varepsilon_j$, estimated from the spread of $\Delta t$, contributes variance $\sigma_j^2$. The total timing variance for a single camera-based timestamp is then

$$\sigma_t^2 = \frac{\Delta T^2}{12} + \sigma_j^2,$$

where $\Delta T$ is the frame interval and $\sigma_j$ the residual synchronization/detection jitter measured empirically. For our setup ($\Delta T \approx 33$ ms) we obtained $\sigma_t \approx 37$ ms. Because gait events (foot-off and foot-strike) are also derived from video frames, their timing relative to the Intan clock is subject to the same uncertainty. However, crucially, the Intan clock, camera triggers, and neural dynamics are not phase-locked, so these errors are effectively random across cycles

##### *Gait cycle duration*

Same-limb gait cycles were defined by two consecutive events at true times $T_0$ and $T_1$, with observed times

$$\hat{T}_k = T_k + \varepsilon_k, \quad \mathrm{Var}(\varepsilon_k) = \sigma_t^2,\ k \in \{0,1\},$$

and true and estimated durations

$$D = T_1 - T_0, \qquad \hat{D} = \hat{T}_1 - \hat{T}_0.$$

Assuming independent errors, the duration error satisfies

$$\mathrm{Var}(\hat{D} - D) = 2\sigma_t^2, \qquad \mathrm{SD}(\hat{D}) = \sqrt{2}\,\sigma_t.$$

Numerically, with $\sigma_t \approx 37$ ms,

$$\mathrm{SD}(\hat{D}) \approx 52 \text{ ms}.$$

When estimating the timing of neural events relative to many cycles, the standard error of the mean scales approximately as

$$\mathrm{SEM}_{\mathrm{cycle}} = \frac{\sqrt{2}\,\sigma_t}{\sqrt{N}} \approx \frac{52 \text{ ms}}{\sqrt{50}} \approx 7.4 \text{ ms},$$

about 1.5% of a $D \approx 500$ ms gait cycle.

##### *Neuron timing relative to cycle onset*

Neuron peak firing times $s_{i,n}$ were expressed relative to the onset of each gait cycle $n$, delimited by $\hat{T}_{0,n}$ and $\hat{T}_{1,n}$. Defining the true and estimated peak times as

$$\tau_{i,n} = s_{i,n} - T_{0,n}, \qquad \hat{\tau}_{i,n} = s_{i,n} - \hat{T}_{0,n} = \tau_{i,n} - \varepsilon_{0,n},$$

gives the single-event timing uncertainty

$$\mathrm{SD}(\hat{\tau}_{i,n}) = \sigma_t \approx 37 \text{ ms},$$

i.e. about 7% of a 500 ms cycle.

##### *Neuron phase within the gait cycle*

For each cycle $n$, delimited by $\hat{T}_{0,n}$ and $\hat{T}_{1,n}$, we defined

$$\hat{D}_n = \hat{T}_{1,n} - \hat{T}_{0,n}, \qquad \hat{\phi}_{i,n} = \frac{s_{i,n} - \hat{T}_{0,n}}{\hat{D}_n}.$$

Treating phase as $f(\tau, D) = \tau/D$ and using first-order error propagation with

$$\frac{\partial f}{\partial \tau} = \frac{1}{D_n}, \qquad \frac{\partial f}{\partial D} = -\frac{\tau_{i,n}}{D_n^2},$$

and

$$\mathrm{Var}(\hat{\tau}_{i,n}) = \sigma_t^2, \quad \mathrm{Var}(\hat{D}_n) = 2\sigma_t^2, \quad \mathrm{Cov}(\hat{\tau}_{i,n}, \hat{D}_n) = \sigma_t^2,$$

gives

$$\mathrm{Var}(\hat{\phi}_{i,n}) \approx \frac{\sigma_t^2}{D_n^2} + \frac{2\tau_{i,n}^2}{D_n^4}\,\sigma_t^2 - 2\,\frac{\tau_{i,n}}{D_n^3}\,\sigma_t^2.$$

For $D_n \approx 500$ ms, $\tau_{i,n} \approx 250$ ms and $\sigma_t \approx 37$ ms,

$$\mathrm{Var}(\hat{\phi}_{i,n}) \approx 0.0021, \qquad \mathrm{SD}(\hat{\phi}_{i,n}) \approx 0.046,$$

i.e. about 5% of the cycle in phase units. Averaging phase across $N$ cycles gives

$$\mathrm{SEM}_{\phi} \approx \frac{\mathrm{SD}(\hat{\phi})}{\sqrt{N}}.$$

Using $\mathrm{SD}(\hat{\phi}) \approx 0.046$ and $N = 50$,

$$\mathrm{SEM}_{\phi} \approx 0.0065, \qquad \mathrm{SEM}_{\mathrm{time}} \approx \mathrm{SEM}_{\phi} D \approx 0.0065 \times 500 \text{ ms} \approx 3.3 \text{ ms},$$

i.e. about 0.7% of the cycle duration. Thus, even though individual cycles are limited by the ∼33 ms frame interval, the means of neural events can still be estimated with substantially finer effective temporal precision. For the parameter values used here, this indicates that neural events can be localized within the gait cycle with an effective precision of only a few milliseconds, corresponding to 2% or less of the step cycle for rats walking are relatively slow speed (2Hz), despite the coarser resolution of the underlying camera frames.

#### Population activity, PCA

For each animal, matrices of time × neuron firing rates were formed for the relevant epochs (locomotion, natural stops, PPN-induced stops, perturbations). A mean-cycle firing matrix was constructed by dividing each gait cycle into a fixed number of phase bins and averaging the rates within each bin across good cycles. Here, an additional stability criterion was applied to select consecutive cycles where the overall unit stability was high to construct the most representative embedding space. This matrix has dimensions (phase bins × neurons).

Principal component analysis (PCA) was applied to the mean-cycle matrix. The first three principal components (PCs) captured most of the cycle-related variance and defined a 3D latent space. The mean locomotor cycle traced a closed loop in this space, which we refer to as the locomotor manifold. Using the same PCA basis, individual cycles, stop epochs, and perturbation trajectories were projected into the same space to compare their geometry and dynamics.

#### Manifold in latent space

To characterize the support of the manifolds, we constructed alpha shapes ('alphaShape.m', in Matlab) on the cloud of latent points corresponding to locomotor and postural states. An alpha shape is a refined version of a convex hull that encloses a cloud of points, while allowing concave boundaries and disregarding sparsely populated regions, thereby capturing the main region in the latent space that the data occupy. All manifolds in latent space (for locomotor, natural stop, PPN-stimulated stops, and post-perturbation) were constructed from subsampled and denoised sets of latent points to obtain robust shapes that reflect typical neural states rather than being dominated by long epochs or outliers. For each behavioral condition, we first identified contiguous time segments that met the relevant criterion and had a minimum duration ($\geq 0.5$ s) to exclude very brief transient events. The time indices of these segments were concatenated, randomly shuffled, and then downsampled by a fixed factor to achieve an approximately uniform sampling density in the 3D latent space. The corresponding latent coordinates (the first three principal components) were cleaned by removing statistical outliers, after which an alpha shape was constructed from the remaining points using a fixed alpha parameter that captured all points. This procedure ensured that all manifolds were defined by dense, representative cores of neural states, making comparisons of geometry and dynamics (for example, flow fields and alignment) across conditions directly interpretable. The overlap between manifolds in the latent space was quantified by comparing the alpha shapes constructed for each condition. For a given pair of manifolds, we identified all latent points that lay inside both alpha shapes and computed the volume of the overlapping region, as well as the fraction of each manifold's volume that was shared. This yielded a symmetric overlap measure that reports, for example, what proportion of the PPN-induced postural manifold is contained within the natural postural manifold, and vise versa, thus capturing how similar or distinct the neural state supports are across behavioral conditions.

#### Flow fields in latent space

To quantify neural dynamics, we computed flow fields in the 3D latent space defined by the first three principal components (**Fig. 3-4**). For each time step, we estimate the instantaneous velocity of the neural state as $v(t) = [y(t+\Delta t) - y(t)]/\Delta t$, where $y(t)$ is the latent state, and $\Delta t$ is a fixed step on the common time grid. The latent space was partitioned into "spatial" bins (i.e., volume bins of the state space), and for each bin $b$ we collected all velocity vectors whose origin lay in that bin, denoted $\{v_{b,i}\}_{i=1}^{N_b}$. The mean local flow in bin $b$ was

$$\bar{v}_b = \frac{1}{N_b} \sum_{i=1}^{N_b} v_{b,i}.$$

Bins with too few samples were discarded. These vectors $\bar{v}_b$ were used to visualize the direction and magnitude of population flow within the locomotor and postural manifolds. Time-shuffled trajectories were used to construct shuffled vector fields as a control for all subsequent analyzes.

#### Flow Alignment

The alignment calculation (**Fig. 3g**) assesses how closely the trajectory during each cycle follows the locally defined flow field, providing a measure of how "organized" the dynamics are within each manifold. For each cycle or stop $k$, we use the velocity vectors $v(t)$ computed at consecutive time steps. The tangent of the trajectory at time $t$ within cycle/stop $k$ is given by the corresponding velocity vector normalized to unit length,

$$\hat{u}_k(t) = \frac{v(t)}{\|v(t)\|}$$

At the same time point, we identify the spatial bin $b_k(t)$ that contains the latent state $y(t)$ and take the mean flow in that bin, $\bar{v}_{b_k(t)}$, normalized as

$$\hat{v}_{b_k(t)} = \frac{\bar{v}_{b_k(t)}}{\|\bar{v}_{b_k(t)}\|}$$

The local alignment between the cycle trajectory and the flow field is defined as the cosine of the angle between these two unit vectors,

$$a_k(t) = \hat{u}_k(t) \cdot \hat{v}_{b_k(t)}$$

A single alignment score for cycle $k$ is then obtained by averaging over all time samples in that cycle/stop,

$$A_k = \frac{1}{L_k} \sum_{t \in \text{cycle } k} a_k(t)$$

where $L_k$ is the number of time points in the cycle. High $A_k$ values indicate that the trajectory closely follows the locally defined flow (organized dynamics), while low $A_k$ values indicate that the trajectory cuts through the flow (disorganized dynamics). By examining the distribution of $A_k$ across the cycles within the locomotor and postural manifolds, we show that the trajectories during locomotion are highly aligned with the flow, consistent with the dynamics of the limit-cycle, whereas the trajectories during stopping are weakly aligned, consistent with fluctuations around fixed-point attractors.

#### Mean flow within manifolds

To compare how strongly the dynamics act in different regions of latent space, we use the same binned flow field as before, where each spatial bin $b$ has an average velocity vector $\bar{v}_b$ that indicates how the neural state tends to move in that local region (**Fig. 3h**). Bins whose centers fall inside a given alpha-shape (for example, the locomotor or postural manifold) are grouped as "inside," and the others as "outside." The mean flow magnitude inside a manifold is then computed as the average length of the flow vectors from all bins inside:

$$\langle \|\bar{v}\| \rangle_{\text{in}} = \frac{1}{|B_{\text{in}}|} \sum_{b \in B_{\text{in}}} \|\bar{v}_b\|.$$

The mean flow magnitude outside is computed in the same way for the set of bins outside the manifold. Intuitively, $\|\bar{v}_b\|$ measures how fast the state tends to move when in the bin $b$, and $\langle \|\bar{v}\| \rangle_{\text{in}}$ or $\langle \|\bar{v}\| \rangle_{\text{out}}$ summarize this "typical speed" inside or outside the manifold. A manifold with relatively strong flow inside compared to outside is compatible with a limit-cycle-like attractor: once the state is on the manifold, it is driven to move along a repeating trajectory. A manifold with weak flow inside but stronger flow in surrounding regions is compatible with a fixed-point-like attractor: once the state is inside, motion is slow and dominated by small fluctuations around stable states, while faster flows outside can push the system toward that region. These inside-outside flow magnitudes are then interpreted together with alignment and Lyapunov analyzes to infer the stability properties of the locomotor and postural manifolds.

#### Cohomology of neural manifolds

Persistent homology was used to characterize the topology of the neural dynamics underlying each manifold (bottom, **Fig. 1g-j** and **3e-f**). For each condition (locomotor, stops), we used the same sets of time points whose latent states defined the alpha shape manifold, so geometric (alpha shape) and topological (cohomology) estimates were based on the same underlying epochs. For each set of time points, population firing rates were projected onto as many principal components as needed to explain at least 95% of the variance, ensuring that the most meaningful high-dimensional structures of the dynamics were retained rather than being compressed into a small fixed number of dimensions. Several circularly shifted copies (0, 50, 100 samples) of these trajectories were then concatenated to build a time-embedded representation, so that each point encoded a short segment of the dynamics rather than an isolated instantaneous state. This procedure preserves the topology of smooth attractors while making loops or fixed points easier to detect from the sampled data.

The resulting high-dimensional signal was downsampled and embedded in three dimensions using a dimensionality reduction method, Uniform Manifold Approximation and Projection (UMAP, cosine metric, 50 neighbors, min_dist 0.15), which approximately preserves local neighborhood relations and thus tends to maintain large-scale topological features such as the presence or absence of loops. Using a tool in topological data analysis that builds a series of shapes (Vietoris–Rips complexes) from a point cloud topological features, like holes and solids, was applied. Persistent homology was computed in these 3D point clouds up to dimension 2 using an open source package of persistent homology for Python, `ripser`[23], in default radius settings, producing barcodes for $H_0$, $H_1$, and $H_2$. The long-lived features of $H_0$ correspond to stable connected components, while the long-lived features of $H_1$ correspond to robust loops. The same pipeline was applied to time-shuffled controls obtained by independently shuffling each principal component, which preserves statistics but destroys temporal structure. To identify robust topological features rather than random fluctuations, we used the persistence diagrams obtained from the shuffled data as an empirical null. Specifically, for each animal and homology dimension, we calculated the 99th percentile of the persistence distribution in the shuffled diagrams and defined a feature in the original data as "robustly persistent" if its persistence exceeded this shuffle-based threshold in the corresponding dimension. The resulting count of such robustly persistent bars in each dimension was used as a statistic reporting the number of topological features whose persistence could not be explained by the shuffled null model. This procedure ensures that, in cases where no structure is present beyond the shuffled baseline, no features are spuriously labeled as persistent. Under locomotor conditions, the diagrams showed a single prominent persistent loop in $H_1$ and no stable $H_2$, consistent with a ring-like attractor, while stop manifolds showed only one robust component in $H_0$ without persistent loops, consistent with a solid region of fixed-point-like states. These topological results align with the flow-based metrics, together indicating coherent rotation on a ring-shaped attractor during locomotion and low-flow fluctuations around stable fixed points during stops, and supporting that the observed topology reflects the genuine high-dimensional attractor rather than an artifact of low-dimensional projection.

#### Stability analysis and Lyapunov exponents

To address the attractor properties of the neural population dynamics, we quantify how stable the dynamics were around each manifold by computing finite-time Lyapunov exponents from nearby trajectories in latent space (**Fig. 4i** and **Extended Data Fig. 9** ). This analysis probes whether two almost identical neural states stay close, slowly drift apart, or rapidly diverge as time evolves, thereby distinguishing stable fixed points, limit-cycle-like behavior, and unstable dynamics. For locomotor epochs, the onsets of gait cycles were used as starting points: pairs of cycle-start indices were selected from cycles with similar durations, so that differences in the exponents were not driven by trivial changes in cycle length or progression speed along the locomotor manifold. For stop-related conditions (voluntary and PPN-induced), cycles are not defined, so trajectories were analyzed over a fixed time window whose length was set by the shortest stable stop in that condition (shortest voluntary stop, shortest stimulation-induced stop), ensuring that all trials were compared over an equivalent duration. For perturbation trials, starting points were chosen 600 ms after the perturbation onset, and trajectories were followed forward up to the shortest available end time across perturbations, so that exponents specifically quantified the stability of the post-perturbation decay back toward the postural manifold over a matched duration.

For a given pair of starting indices $(i, j)$, the corresponding segments of the trajectory in the first three principal components were extracted over the chosen duration $T$. The relationship between the initial distance $D_0$ and the distance $D_k$ after a discrete time lag $t_k$ was modeled as follows:

$$D_k \approx D_0 \exp(\lambda t_k),$$

where $\lambda$ is the finite-time Lyapunov exponent. The initial Euclidean distance

$$D_0 = \|x_i(0) - x_j(0)\|$$

The distance was computed, and at each discrete time step $t_k$ within the window, the distance

$$D_k = \|x_i(t_k) - x_j(t_k)\|$$

was evaluated. A finite-time Lyapunov exponent for that pair was then estimated in discrete form as:

$$\lambda_{ij} \approx \frac{1}{K} \log\left(\frac{D_K}{D_0}\right),$$

where $K$ is the number of valid time steps in the window, providing a discrete approximation to the average growth (or decay) per-step of small perturbations in latent space. All pairwise exponents $\lambda_{ij}$ were assembled into a symmetric matrix, resulting in a spectrum of local stability values across the manifold. The mean of this spectrum was used as a summary measure of typical stability under each condition: a negative mean indicated that nearby trajectories generally converged (stable, fixed-point-like dynamics), a mean near zero indicated neutrally stable, limit-cycle-like dynamics on the locomotor manifold, and a positive mean would indicate overall instability. This procedure was applied to (i) locomotor cycles ("AllCycles"), to assess the stability of the locomotor manifold; (ii) voluntary and PPN-induced stops, to quantify convergence toward postural fixed-point manifolds; and (iii) perturbation trials, to measure how quickly deviations caused by transient treadmill jerks relaxed back to the postural manifold.

#### Classification of neuronal modulation across conditions

To characterize how individual neurons contributed to locomotor and postural states, mean firing rates were calculated for each neuron during locomotion and during sustained stopping (voluntary and PPN-induced). For each neuron, a movement modulation index was defined as the ratio between its firing rate during locomotion and during stopping, providing a single scalar that summarizes how strongly the neuron was biased toward movement versus posture. Using this index and a fixed threshold $p_{\text{rc}} = 0.1$, neurons were assigned to four modulation classes. Silenced neurons had a modulation index $< 0.1$, indicating that their firing almost vanished during stopping relative to locomotion; The decreased neurons had an index between 0.1 and 1, reflecting a moderate reduction during stops. The increased neurons had an index $> 1$ but with an inverse index (“stop”/“locomotion”) $\geq 0.1$, indicating a moderate increase in firing during stops. The neurons that appeared had an inverse index $< 0.1$, which means that they were almost inactive during locomotion, but became strongly active during stopping. The distribution of these classes was quantified across animals, revealing that the majority of neurons decreased their firing or were silenced during stops, while smaller fractions increased or appeared, consistent with a broad suppression of locomotor drive and recruitment of a more specialized postural population.

##### Probe reconstruction and laminar assignment

To localize recorded units within the spinal cord, Neuropixels probe geometries were registered in animal-specific MRI segmentation of the lumbar cord generated using open source software (3D Slicer, `https://www.slicer.org/`). For each rat, white matter, gray matter, dorsal horn, intermediate zone, and ventral horn were segmented as 3D meshes (STL files). The track left by the implanted probe shaft was segmented in the same 3D Slicer project and exported as an STL in the same MRI space (**Fig. 2b**). The Neuropixels layout map (the fixed 3D coordinates of all recording sites) was rigidly aligned to the segmented track using an orthogonal Procrustes procedure. To enforce the one-to-one correspondence required by this method, a set of points equal in number to the Neuropixels channels was randomly sampled from the segmented track, and a rotation and translation were estimated via singular value decomposition to minimize the mean squared distance between the corresponding points. Then a final small translation was applied to align the tip of the layout map with the ventral part of the segmented track.

Alpha shapes were constructed around the segmentations of the dorsal horn, intermediate and ventral horn, and the aligned channel coordinates were tested for membership in each region. The channels were labeled dorsal, intermediate, or ventral depending on which alpha shape they fell into (with intermediate taking precedence in overlapping zones), and single units were assigned to these laminar groups based on the channel of maximal waveform amplitude, providing a laminar position for each neuron.

#### Supplementary Videos

**Supplementary Video 1** Activity of the neuronal population in the lumbar spinal cord during walking and stopping (PPN-induced, blue regions). Real time unit activity spatially distributed on the Neuropixels shank (Left, down is ventral). Video recording of the rat (R174) while walking volitionally (top middle). Top right, population activity in two-dimensional space using the first two principal components of the neuronal firing rates. Bottom, raster of the population activity (sorted by phase). Note that the sequential activity seems to cover all phases (rotational dynamics). The Blue shaded regions indicate optogenetic stimulation of the brainstem nucleus (PPN), which induces stopping. Below the raster are shown the kinematics (angles of ankle-paw, $\beta_{AP}$, paw-snout, $\beta_{PS}$, and Tail-ankle, $\beta_{TA}$, see "Kinematics angles" and **Extended Data Fig. 6**).

**Supplementary Video 2** Neuronal population activity in the lumbar spinal cord visualized in neural state space during a walk-to-stop transition generated by PPN stimulation. Two experimental data sets shown (rat 173, n=20 trials and rat 174, n=25 trials). Next, the model simulation of a walk-to-stop transition is accomplished by modulation of specific cell types that hence change the projectome, for instance by inhibiting the V1 interneurons or exciting the V2b. The locomotor and postural manifolds are indicated. During the manipulation (PPN-stimulation or V1/V2b-modulation) the trajectories turn magenta.

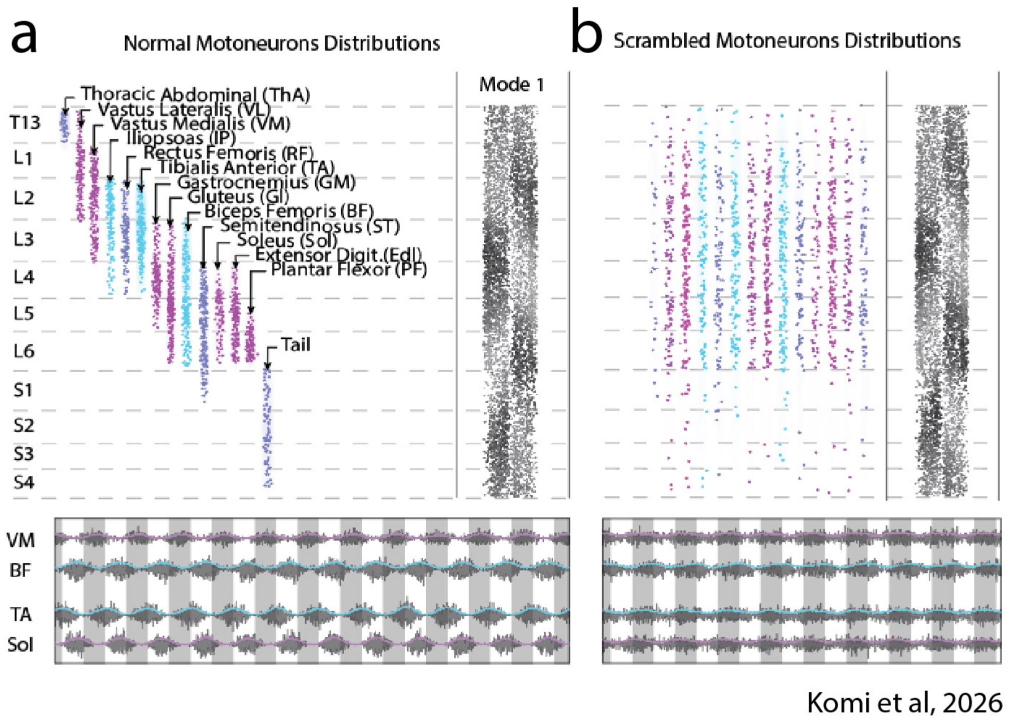


**Extended Data Fig. 1.** | **Appropriate sequential activation of muscles can be achieved by the spatial organization of motoneurons and a traveling wave of neuronal activity that locally activates motoneurons. a**, top left: Experimentally derived spatial organization of motoneurons pools, color coded according to flexor- (cyan) and extentor-types (magenta) for various muscles in the rostrocaudal directions in the rodent (mouse). Non-flexor/extensor motoneuron pools are shown in addition (blue). Top right, a model simulation of a traveling wave of neuronal activity in the longitudal direction. Bottom: The resultant nerve activities of four sample muscles as a result of the wave propagation. This is similar to experimentally observed flexor/extensor activites[1]. **b**, When scrambling the position of the motoneuron pools, the activity pattern of the muscles falls apart (bottom). Motoneuron positions are reproduced with permission from previous reports[39,40]. Model data is reproduced with permission[1].

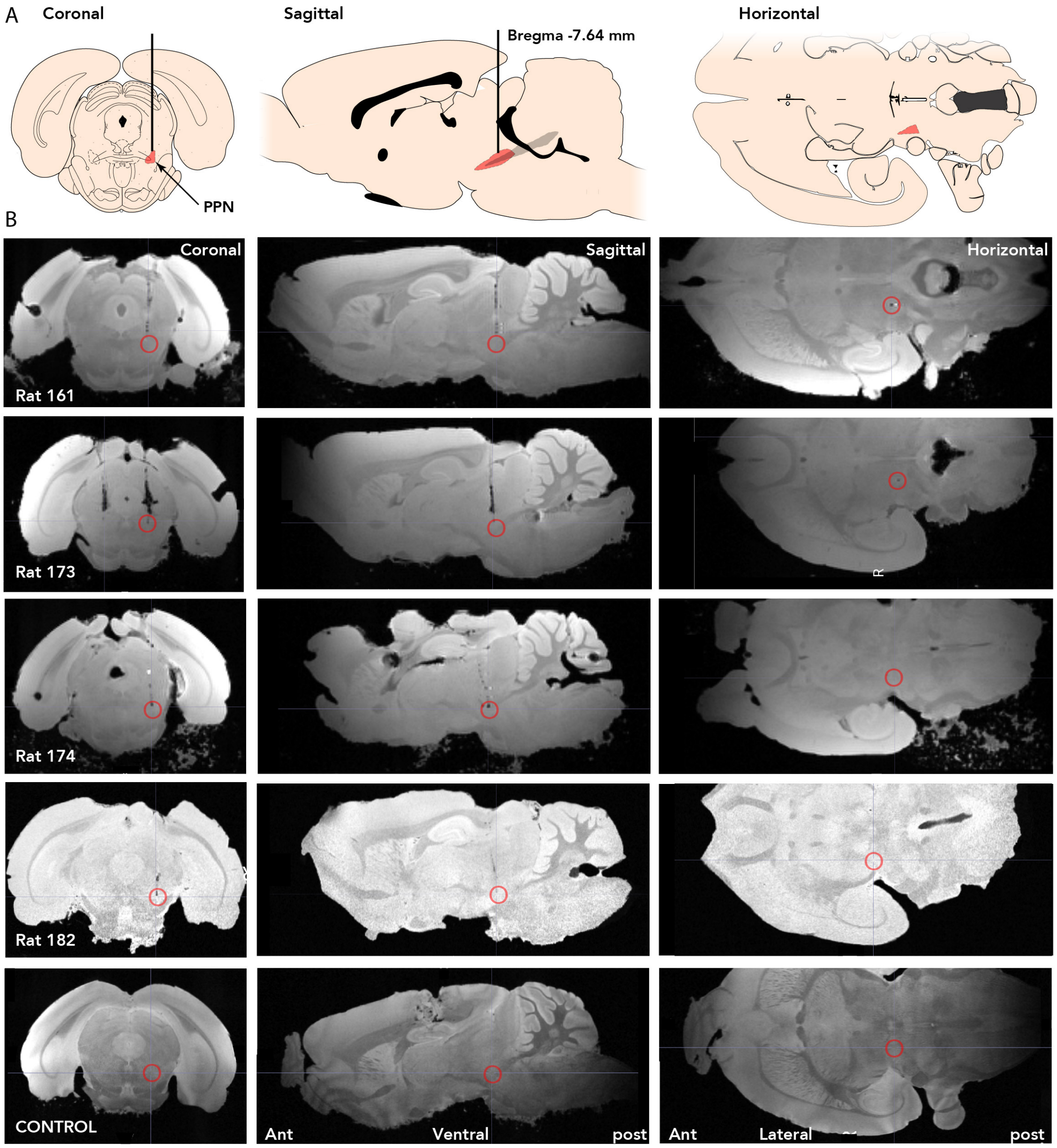


**Extended Data Fig. 2.** | **Location of optical probe across a subset of animals. A**, Illustration of the location of the pedunculopontine nucleus (PPN, indicated red) in coronal (left), sagittal (middle) and horizontal sections (right). **B**, Post mortem brain scannings (MRI) of a subset of animals with the location of the optical probe indicated (red circle) in coronal (left), sagittal (middle) and horizontal (right) sections. Bottom panel shows a control scan, i.e. a scan of a rat brain without an optical probe implanted in PPN. Sections in **a** reproduced with permission[41].

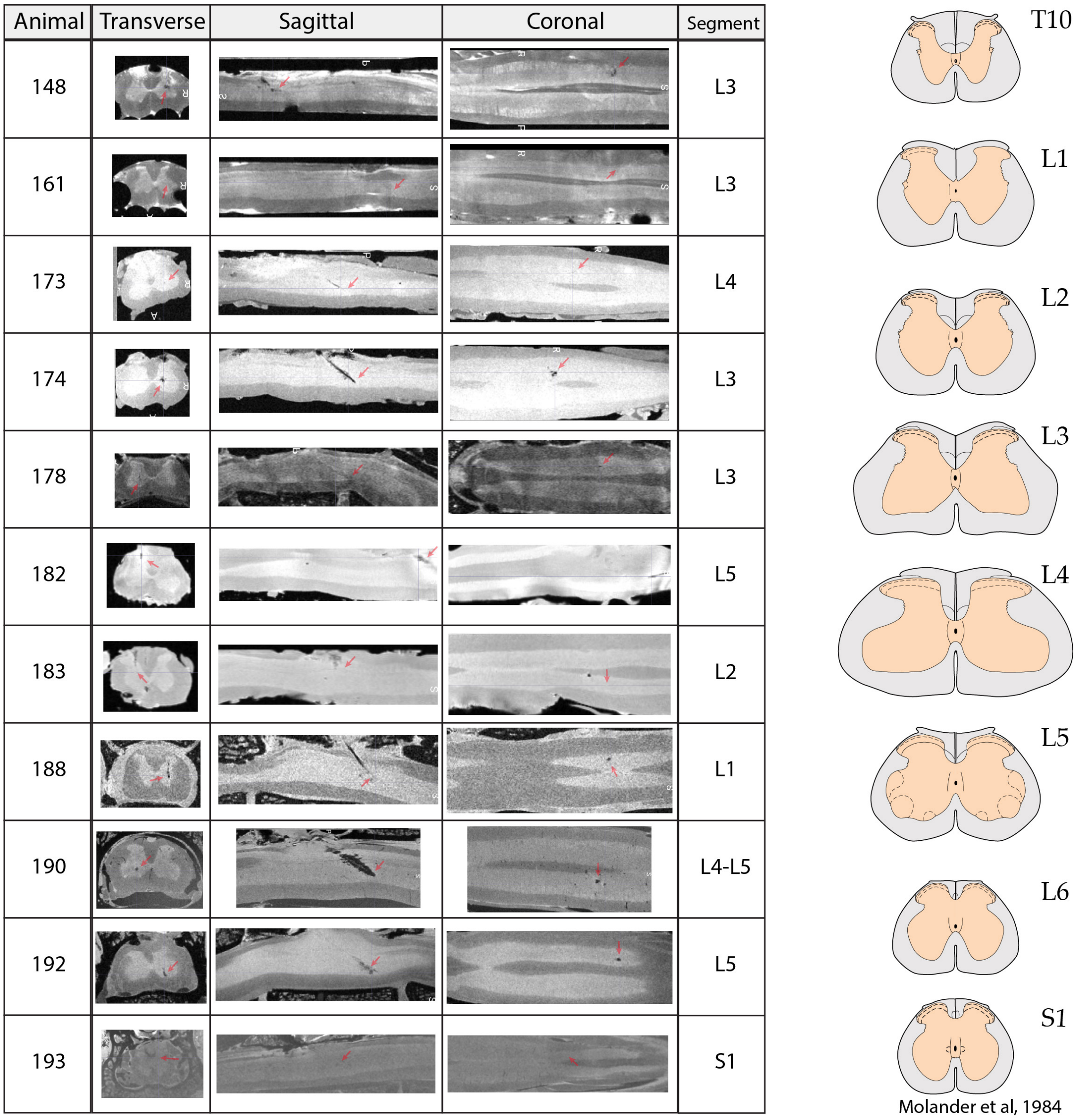


**Extended Data Fig. 3.** | **Location of Neuropixels probe across animals. Left**, Table containing structural MRI image of the spinal cord where the Neuropixels shank was in in the tissue (red arrows), in transvers (second left), sagittal (middle) and coronal sections (second right). The identified spinal segment (right column) is based on visual shape-matching to the anatomical atlas of the rat spinal cord (right, relevant segments shown). Spinal reference sections reproduced with permission[42].

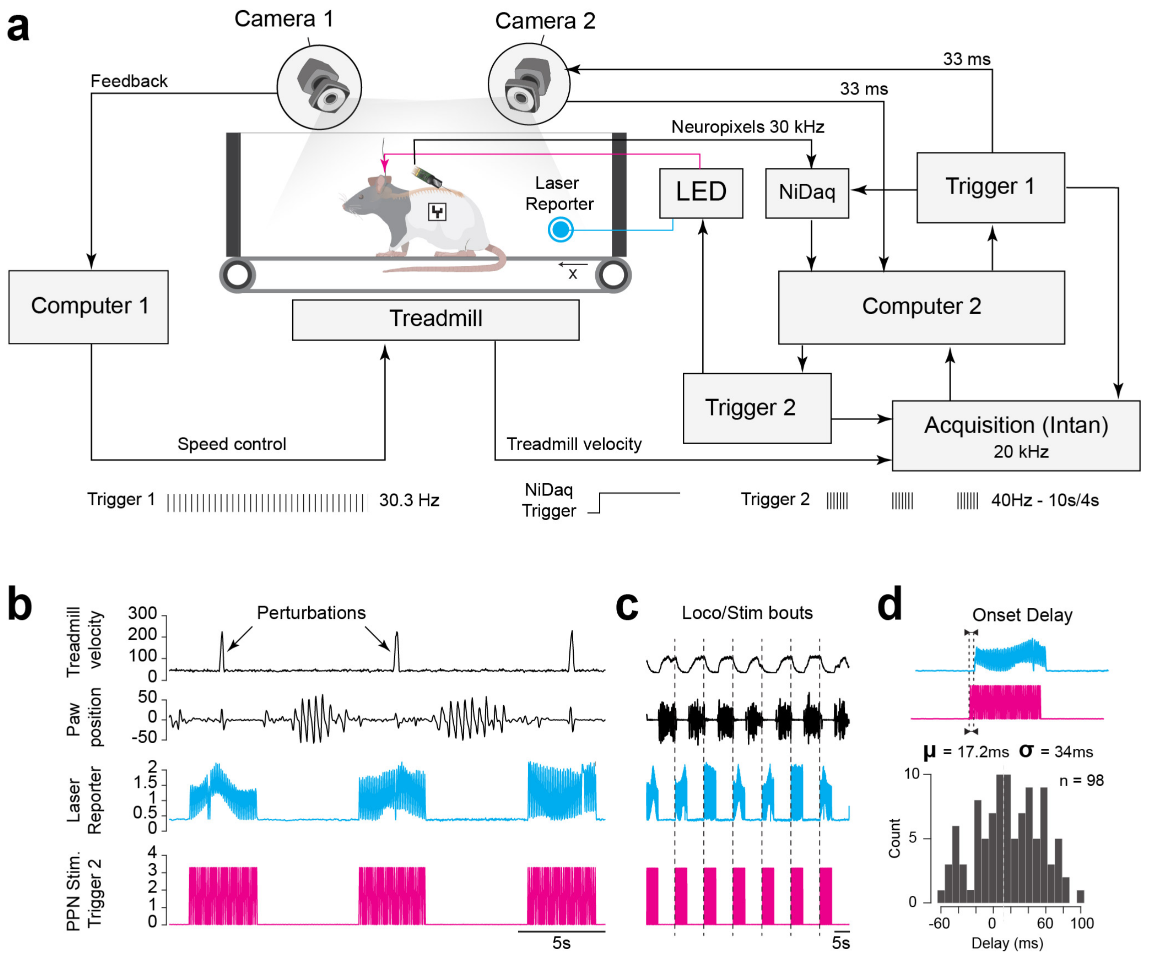


**Extended Data Fig. 4.** | **Experimental setup and assessment of sampling limits. a**, The behavioral experiment consisted of a computer controlled treadmill, where a fiducial marker on the rat is identified by a camera (camera 1) and a computer (computer 1) is adjusting the velocity of the treadmill belt. A second camera (camera 2) is recording the rats movement (at 33 ms intervals) for kinematics, which is recorded on the acquisition computer (computer 2). A trial is initiated by computer 2 through a trigger system (trigger 1, at interval of 30.3 Hz) that synchronizes onset of Neuropixels recording (through the national data acquisition card, "NiDaq"), and pulse triggering the camera 2 (triggering signals indicated at the bottom). The triggering is recorded at an external acquisition board (Intan tech, at 20 KHz) and this is recorded on Computer 2. The computer 2 also triggers the optogenetic stimulation of PPN (trigger 2 and LED), which is recorded by the acquisition board. The treadmill belt velocity is also recorded by the acquisition board, and the perturbation is controlled by manual initiation once per PPN stimulation, which is also recorded by the acquisition board and stored on computer 2. **b**, Sample recording with the belt velocity (top, arb. units), paw position extracted from markerless pose-estimation using the video recording (top second), the PPN stimulation events extracted from the laser reporter from the video recording for verification of alignment (cyan, second from bottom), and the trigger signal for the PPN LED stimulation (magenta, bottom). **c**, Same arrangement as in **b**, but with longer timescale and without perturbation. **d**, Overlay of video-extracted PPN stimulation (cyan, top) and the trigger signal (magenta), as control for correct timing of signals. The video sampling using ∼30 Hz frame rate, occasional frame-dropping and duplication due to buffering issues, introduced uncertainty in the timing of pose signals, which is assessed in the histogram (bottom). The mean bias was 17.2 ms, which could be compensated for and standard deviation of 34 ms.

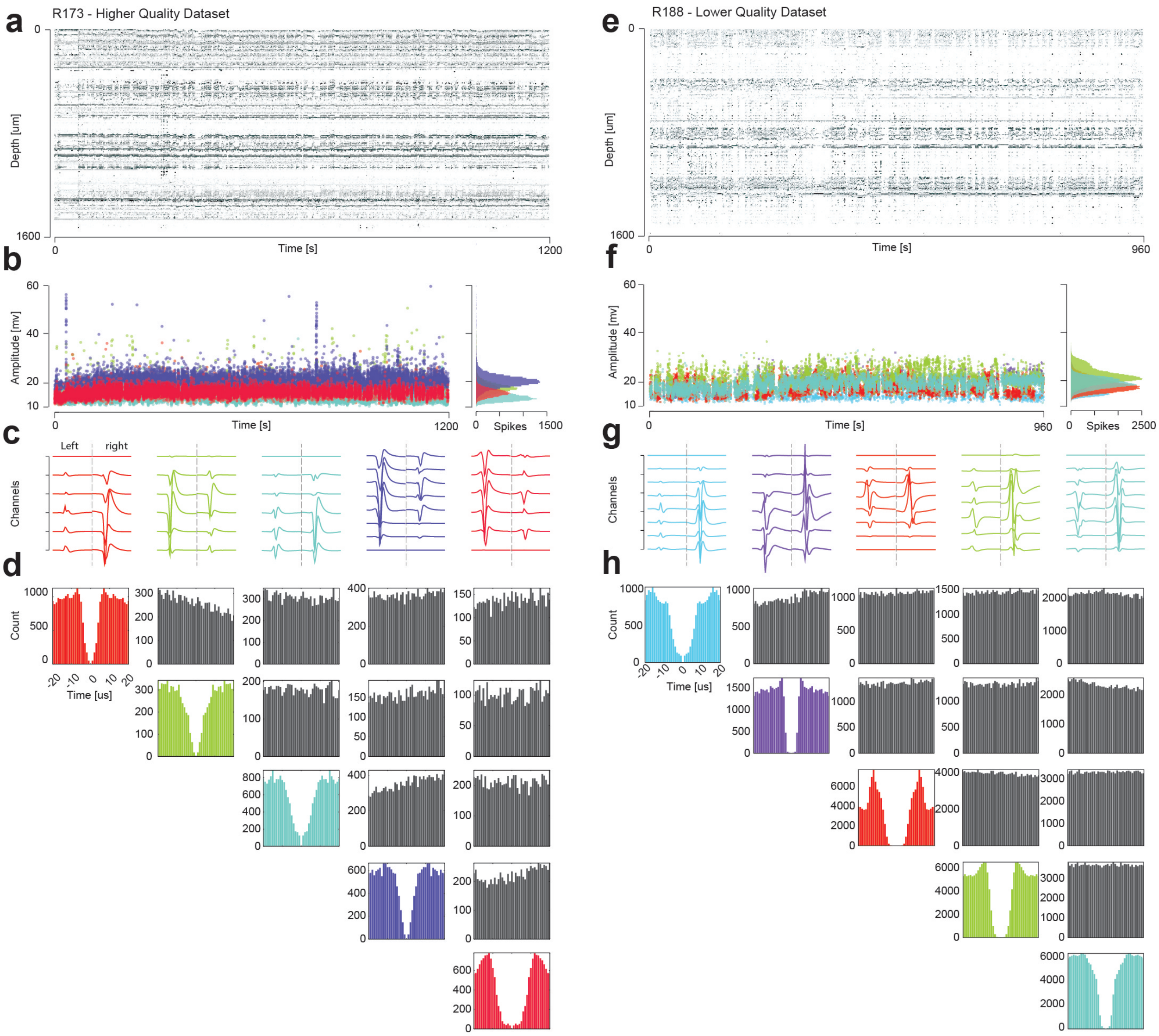


**Extended Data Fig. 5.** | **Stability of Neuropixels recording in spinal cord shown for two sample data sets during walking.** **a**, Drift map of a high quality dataset (Rat 173) recording over 1200 second. Amplitude of the sorted units (469 units) are shown (normalized gray-scale) at different depths (y-axis). **b**, Stability over time can be evaluated via the spike waveform amplitudes for five selected units (colors) over time. Distributions of amplitudes (right). **c**, The associated five sorted sample units from the Neuropixels channel map. Note that the map has a right and left side (divided by broken line), such that the unit appears twice on various channels. **d**, Auto-correlograms (colored diagonal) for the units demonstrate the quality of sorting. Cross-correlograms (gray off-diagonal) verify any cross-talk between channels or possible relationships. **e-h**, Same arrangement as a-d, but for a different data set of lower quality (93 units). Both datasets have been corrected for drift using the Kilosort4 algorithm[37]. Residual drifts are 1.6 $\mu m$ (a) and 3.8$\mu m$ (e).

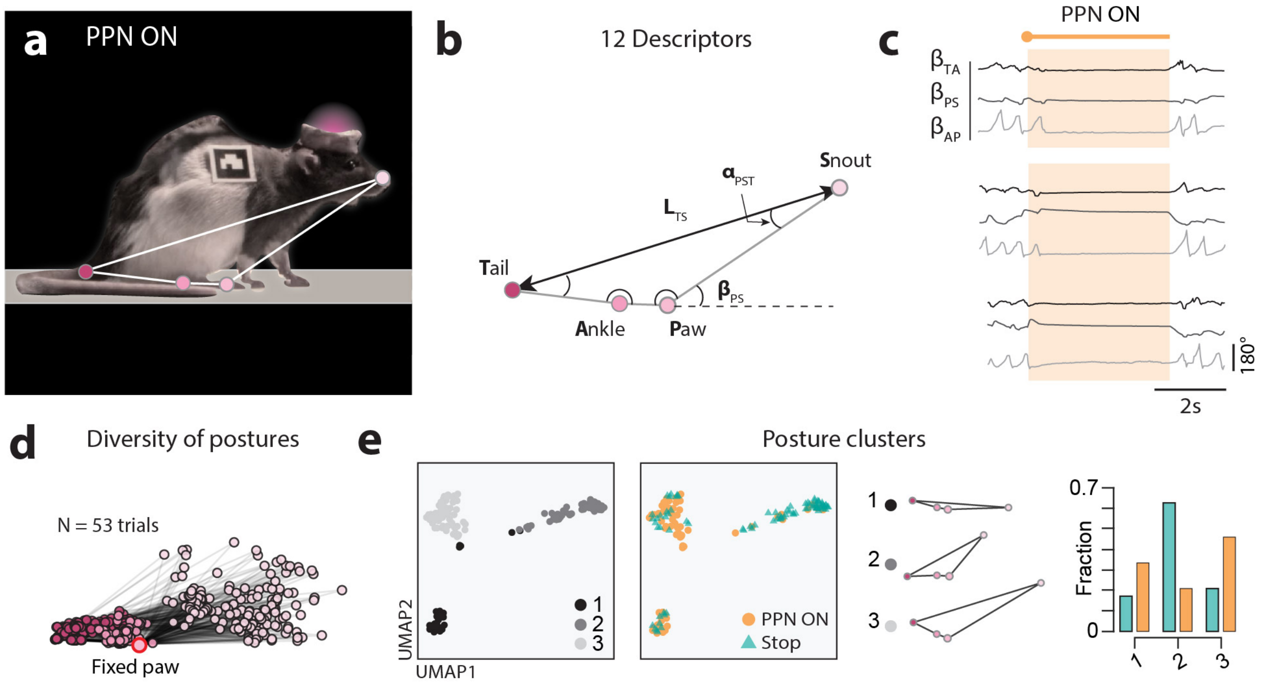


**Extended Data Fig. 6.** | **Diversity of posture during natural- and PPN-induced stops. a**, Sample still frame from videography to illustrate the bodily points (colors) and lines between them (white overlay) used for markerless pose estimation. **b**, The body points used were the base of tail, ankle, hind paw and snout, and the line between then and angles between them formed a total of 12 descriptors. **c**, The kinematics descriptors used in our study were: angles of ankle-paw, $\beta_{AP}$, paw-snout, $\beta_{PS}$, and tail-ankle, $\beta_{TA}$, see "Kinematics angles". Here shown 3 trials of walk-to-stop transitions induced by PPN-stimulation ("PPN-on", yellow shading). **d**, Overlay of a diversity of poses during stopping (n=53 trials) fixed on the paw. **e**, Clustering of poses using UMAP identifies three clusters (left). Both the natural (green) and PPN-induced stopping (orange) had points in all three posture clusters (second left). The mean shapes for the clusters indicates their different postures (second right). The fraction of trials for natural (green) and PPN-induced stopping (orange) are substantial in all three postures.

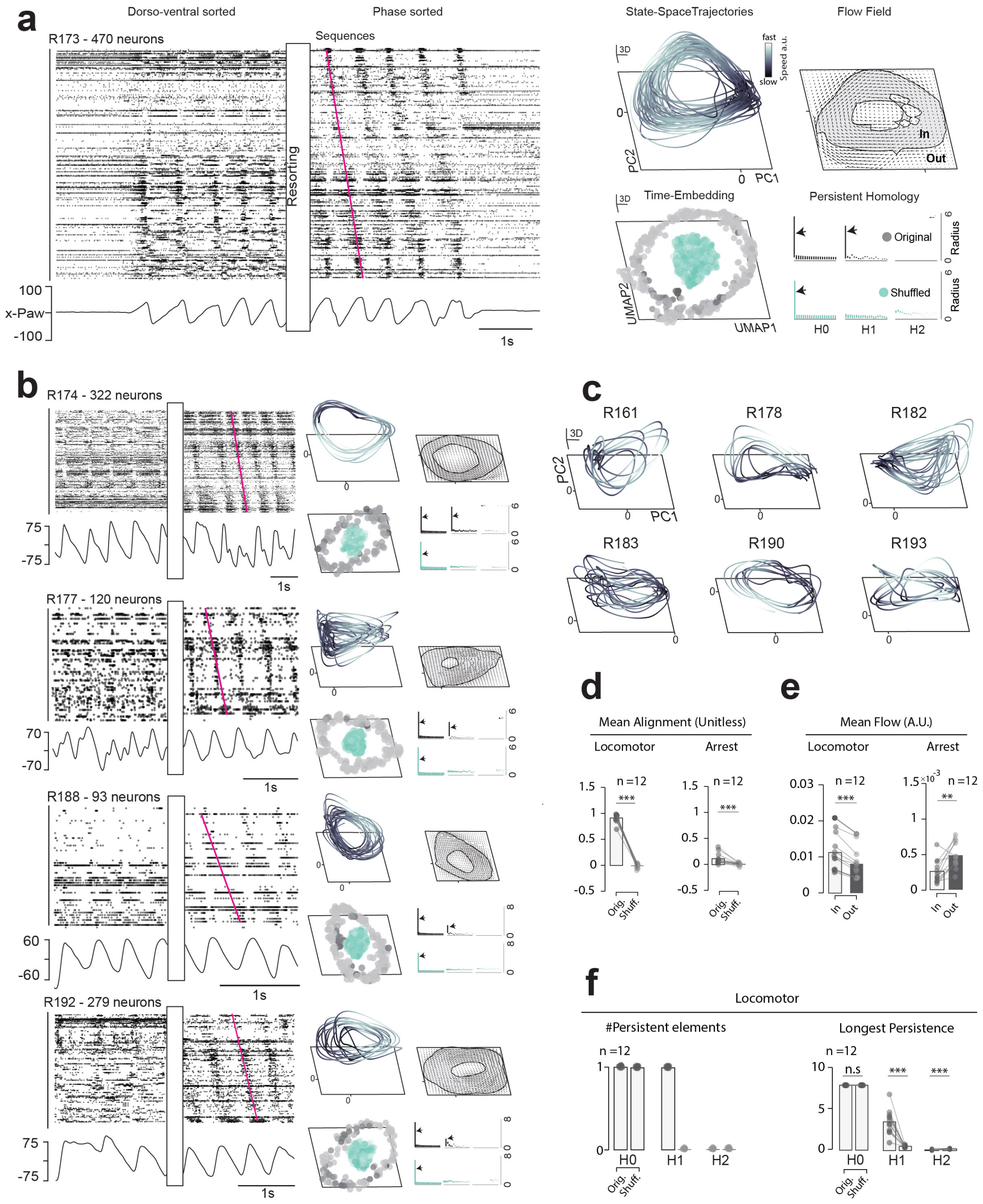


**Extended Data Fig. 7.** | **Rotational population dynamics, manifolds and flow fields across animals.** **a**, Spinal population activity plotted as rasters in dorsoventral sorting along the shank (left) and resorted according to preferred locomotor phase (right). The population dynamics has a sequential progression covering all phases (rotational dynamics, red line) also apparent as a low dimensional circular movement in neural state space (first three principal components, top right). Flow field illustrates the attractive circular motion (far top right) and the topological measures (UMAP, right bottom) demonstrate that the manifold is a solid ring structure (two peaks, arrows, in persistent homology). Time-shuffling of the data points destroyed the ring-topology manifold structure (green points). Homology was performed on the UMAP and time-embedded data. Sample in a is rat 173. **b**, Four highligted other sample animal data sets shown with same structure as in a (Rat174, 177, 188, 192, respectively). **c**, Neural state space for the remaining data sets (rat 161, 178, 182, 183, 190 and 193, respectively). **d**, Mean alignment for all animals during locomotion (left) and stop (arrest, right, n=12). **e**, Mean flow for all animals during locomotion (left) and stop (arrest, right, n=12). **f**, Persistent homology groups elements across population (n=12). Wilcoxon signed-rank test, **: $p < 0.01$., ***: $p < 0.001$.

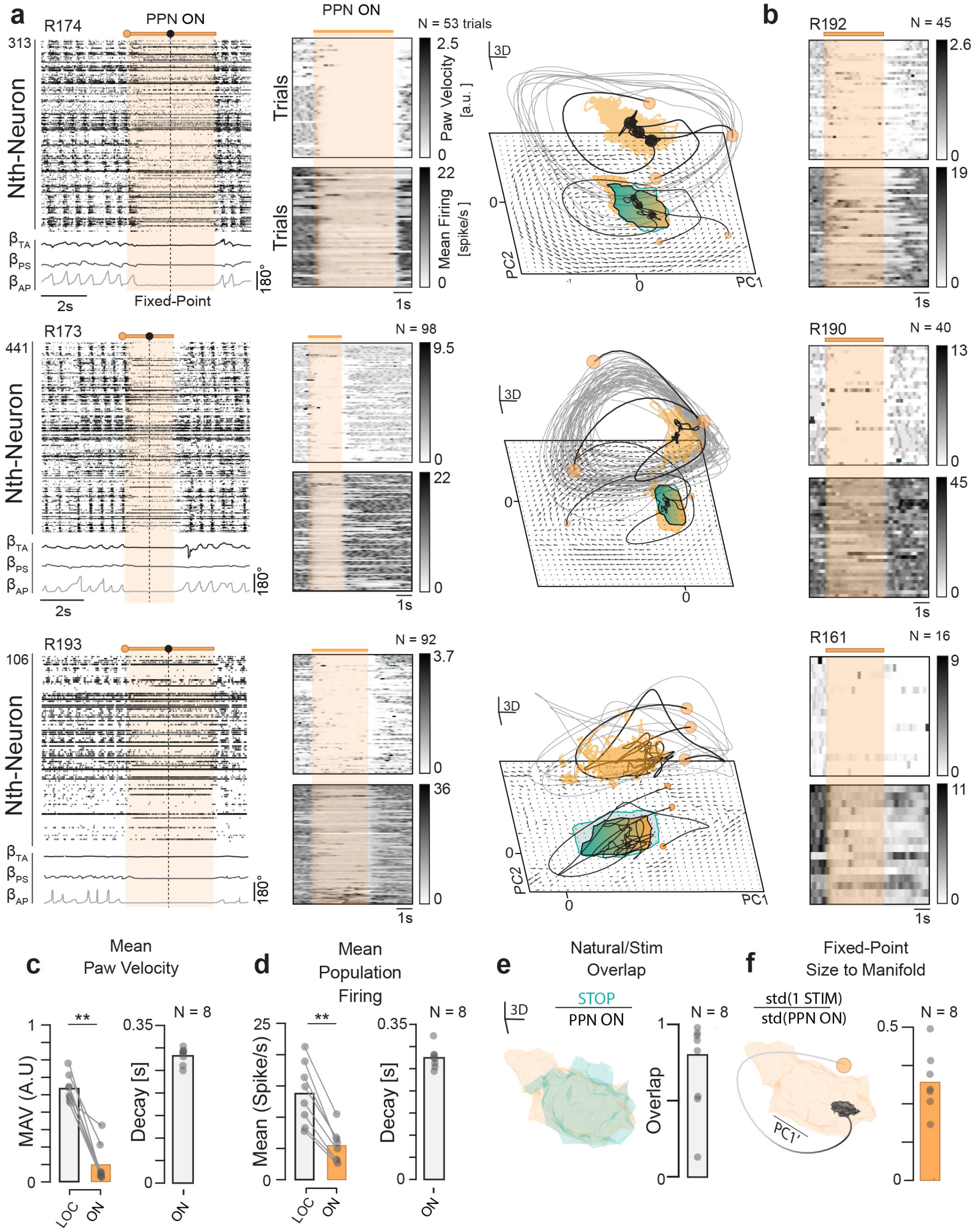


**Extended Data Fig. 8.** | **Induced stopping and postural manifold across animals. a**, Spinal population activity during walk-to-stop transitions across animals (PPN-stimulation: yellow shaded, broken line: point of steady-state assumption, animal R174, R173 and R193). Paw kinematics (bottom). Middle column: Paw velocity and mean firing across trials (n=53/top, n=98/middle, n=92/bottom). Right column: In neural state space, walk-to-stop transition show convergence from locomotor manifold (gray trajectory) to postural manifold (yellow region, PC1-3). **b**, same as middle column in a, except for more animals (R192/n=45, R190/n=40, R161/n=16 trials). **c-d**, Mean absolute paw velocity (MAV), mean firing, and their decay time ("Decay"). **e**, Postural manifold for natural stopping (green) and PPN-induced stopping (yellow). Bar graph: overlap in PC1 across animals (n=8). **f**, Size of fixed point relative to size of postural manifold (posture during PPN-induced stopping, n=8). d, e: Wilcoxon signed-rank test, **: $p < 0.01$.

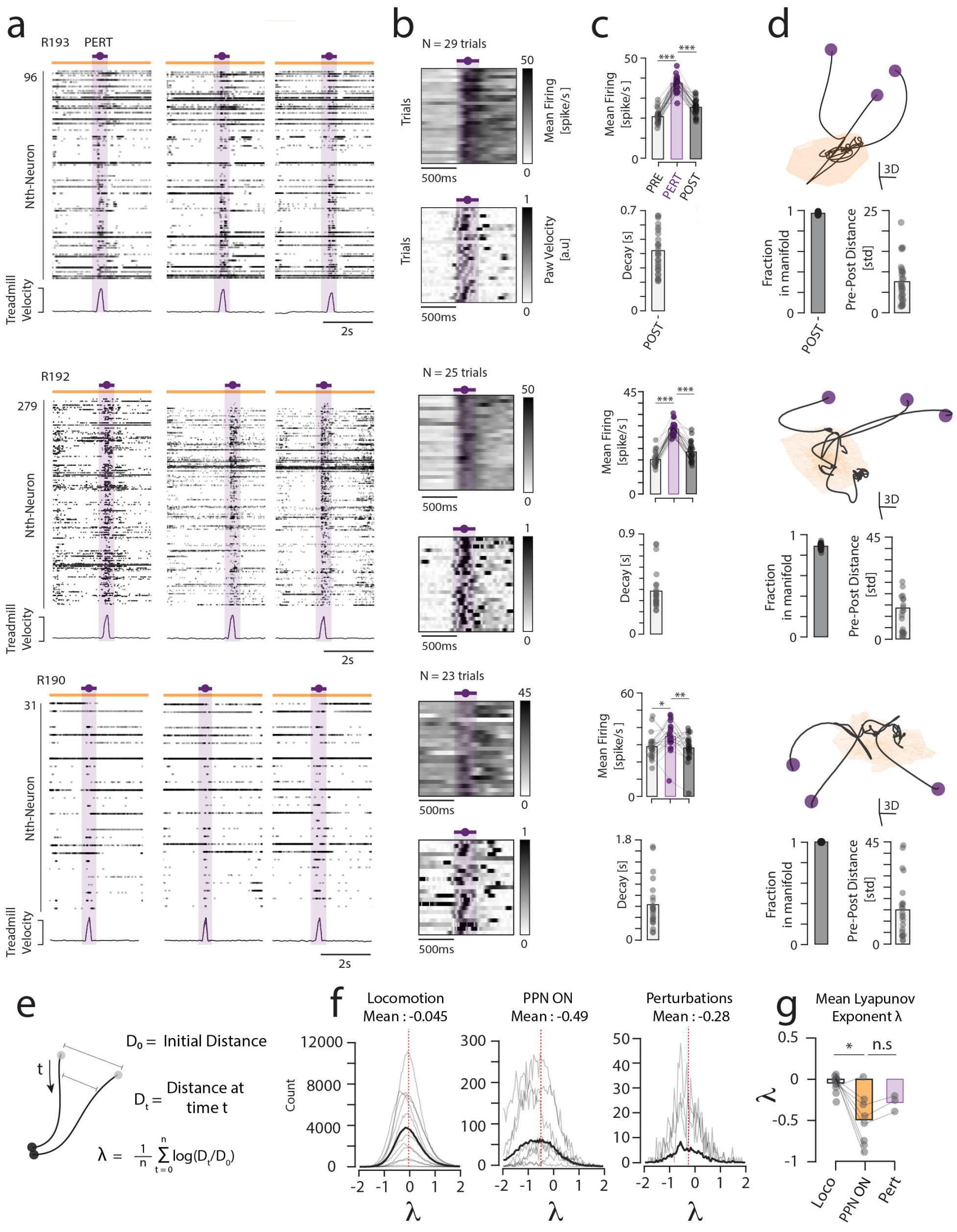


**Extended Data Fig. 9.** | **Perturbation during PPN-induced stopping and postural manifold. a**, Spinal population activity shown as raster plot during PPN-induced stopping: 3 animals (column) each having 3 trials (rows). Half-way through the stop, the limb position is perturbed forward (purple shades, limbs do not return to initial position) with a small forward jerk of the treadmill (velocity at bottom). Animals shown: R193 (top), R192 (middle) and R190 (bottom). **b**, Mean population firing rate before, during and after the perturbation (top) and associated paw velocity (bottom) across animals and trials (R193/n=29, R192/n=25, R190/n=23). **c**, Mean population firing rate before ("PRE"), during ("PERT") and after ("POST") perturbation (top). Bottom: the trajectory time for decay back to postural manifold ("Decay"). **d**, Trajectory in neural state space decay back to postural manifold (shaded region) in neural state space. Fraction of points within the manifold after perturbation (bottom left) and the distance between fixed points of before and after perturbation in standard deviations (bottom right). **e**, Rate of convergence of neighboring trajectories in neural state space is quantified using the exponential decay of inter-trajectory distance over time (i.e. the Lyapunov exponent, $\lambda$). **f**, Distributions of $\lambda$, during three conditions: locomotion, PPN-induced walk-to-stop, and natural walk-to-stop. **g**, the mean $\lambda$ is close to zero during locomotion, which indicates stable limit cycle rotation, whereas the walk-to-stop transition has strong convergence (negative $\lambda$). Wilcoxon signed-rank test, *: $p < 0.05$, **: $p < 0.01$, ***: $p < 0.001$.

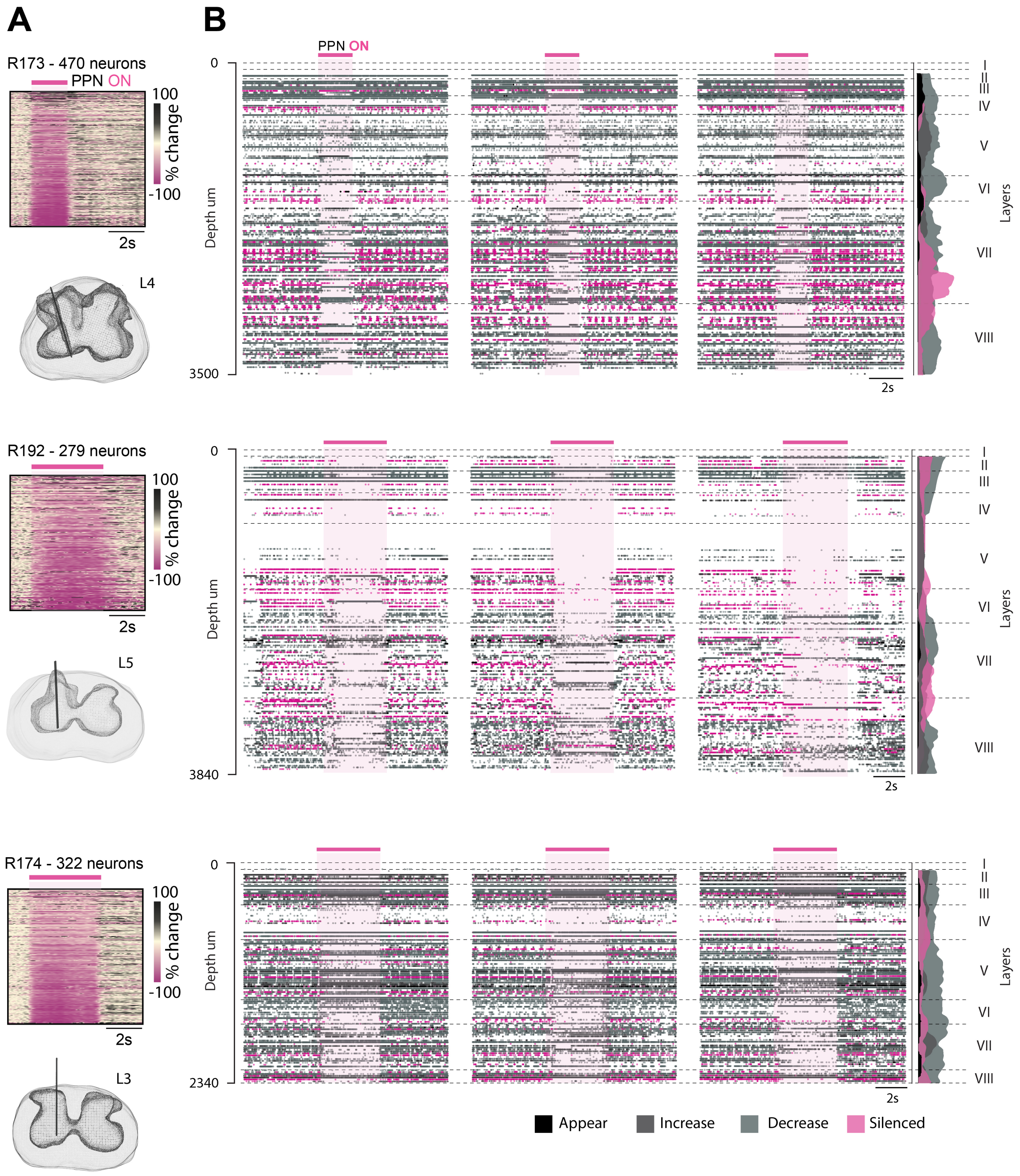


**Extended Data Fig. 10.** | **Neuronal population response during walk-to-stop transition induced by PPN stimulation. a**, Color coded sample firing rates of neurons before, during and after PPN-induced stops (indicated, magenta line) ordered according to firing increase during stop (y-axis). Most neurons have a decrease in firing rate (magenta), although some increase (black). Three animals shown (R173 top/ N=470 neurons, R192 middle/ N=279 neurons, R174 bottom/ N=322 neurons). Bottom: The Neuropixels probe location in transverse spinal sections reconstructed from MRI scans. **b**, Sample rasters of population spiking (three trials for each animal) with the silenced group highlighted (purple). Neuronal location from spinal surface indicated on y-axis ($\mu$m). Rexed layers indicated (right) and the spatial distributions of cellular categories according to response (Appear, increase, reduce and silenced during stops, indicated bottom).

#### References


1. Komi, S. *et al.* Spatial and network principles behind neural generation of locomotion, DOI: 10.1101/2024.10.03.616472 (2024).
2. Wandler, F. D., Lemberger, B. K., McLean, D. L. & Murray, J. M. Coordinated spinal locomotor network dynamics emerge from cell-type-specific connectivity patterns, DOI: 10.7554/eLife.106658.1 (2025).
3. Pazzaglia, A. *et al.* Balancing central control and sensory feedback produces adaptable and robust locomotor patterns in a spiking, neuromechanical model of the salamander spinal cord. *PLOS Comput. Biol.* **21**, e1012101, DOI: 10.1371/journal.pcbi.1012101 (2025).
4. Lindén, H., Petersen, P. C., Vestergaard, M. & Berg, R. W. Movement is governed by rotational neural dynamics in spinal motor networks. *Nature* **610**, 526–531, DOI: 10.1038/s41586-022-05293-w (2022).
5. Wimalasena, L. N., Pandarinath, C. & AuYong, N. Spinal interneuron population dynamics underlying flexible pattern generation. *Nat. Commun.* **16**, 9634, DOI: 10.1038/s41467-025-64629-y (2025).
6. Bush, N. E. & Ramirez, J.-M. Latent neural population dynamics underlying breathing, opioid-induced respiratory depression and gasping. *Nat. Neurosci.* **27**, 259–271, DOI: 10.1038/s41593-023-01520-3 (2024).
7. Bush, N. E., Oliveira, L. M., Glovak, Z. T. & Ramirez, J.-M. An inspiration-off attractor supports the robust and flexible control of breathing, DOI: 10.1101/2025.09.23.678177 (2025).
8. Dale, N. Coordinated Motor Activity in Simulated Spinal Networks Emerges from Simple Biologically Plausible Rules of Connectivity. *J. Comput. Neurosci.* **14**, 55–70, DOI: 10.1023/A:1021176301776 (2003).
9. Sengupta, M., Daliparthi, V., Roussel, Y., Bui, T. V. & Bagnall, M. W. Spinal V1 neurons inhibit motor targets locally and sensory targets distally. *Curr. Biol.* **31**, 3820–3833, DOI: 10.1016/j.cub.2021.06.053 (2021).
10. Sweeney, L. B. *et al.* Origin and Segmental Diversity of Spinal Inhibitory Interneurons. *Neuron* **97**, 341–355, DOI: 10.1016/j.neuron.2017.12.029 (2018).
11. Fetcho, J. R. & McLean, D. L. Some principles of organization of spinal neurons underlying locomotion in zebrafish and their implications. *Ann NY Acad Sci* **1198**, 94–104, DOI: 10.1111/j.1749-6632.2010.05539.x (2010).
12. Liu, Y., Hasegawa, E., Nose, A., Zwart, M. F. & Kohsaka, H. Synchronous multi-segmental activity between metachronal waves controls locomotion speed in Drosophila larvae. *eLife* **12**, DOI: 10.7554/eLife.83328 (2023).
13. Saltiel, P., d'Avella, A., Wyler-Duda, K. & Bizzi, E. Synergy temporal sequences and topography in the spinal cord: evidence for a traveling wave in frog locomotion. *Brain Struct. Funct.* **221**, 3869–3890, DOI: 10.1007/s00429-015-1133-5 (2016).
14. Ryczko, D. *et al.* Flexibility of the axial central pattern generator network for locomotion in the salamander. *J. Neurophysiol.* **113**, 1921–1940, DOI: 10.1152/jn.00894.2014 (2015).
15. Falgairolle, M. & Cazalets, J. Metachronal coupling between spinal neuronal networks during locomotor activity in newborn rat. *The J. Physiol.* **580**, 87–102, DOI: 10.1113/jphysiol.2006.115709 (2007).
16. Cazalets, J. R., Sqalli-Houssaini, Y. & Clarac, F. Activation of the central pattern generators for locomotion by serotonin and excitatory amino acids in neonatal rat. *The J. Physiol.* **455**, 187–204, DOI: 10.1113/jphysiol.1992.sp019296 (1992).
17. Bonnot, A., Whelan, P. J., Mentis, G. Z. & O'Donovan, M. J. Locomotor-like activity generated by the neonatal mouse spinal cord. *Brain Res. Rev.* **40**, 141–151, DOI: 10.1016/S0165-0173(02)00197-2 (2002).
18. Machado, T. A., Pnevmatikakis, E., Paninski, L., Jessell, T. M. & Miri, A. Primacy of Flexor Locomotor Pattern Revealed by Ancestral Reversion of Motor Neuron Identity. *Cell* **162**, 338–350, DOI: 10.1016/j.cell.2015.06.036 (2015).
19. Cazalets, J. Metachronal propagation of motoneurone burst activation in isolated spinal cord of newborn rat. *The J. Physiol.* **568**, 583–597, DOI: 10.1113/jphysiol.2005.086850 (2005).
20. Pérez, T. *et al.* An Intersegmental Neuronal Architecture for Spinal Wave Propagation under Deletions. *The J. Neurosci.* **29**, 10254–10263, DOI: 10.1523/JNEUROSCI.1737-09.2009 (2009).
21. Cuellar, C. A. *et al.* Propagation of Sinusoidal Electrical Waves along the Spinal Cord during a Fictive Motor Task. *J. Neurosci.* **29**, 798–810, DOI: 10.1523/JNEUROSCI.3408-08.2009 (2009).
22. Auyong, N., Ollivier-lanvin, K. & Lemay, M. A. Preferred locomotor phase of activity of lumbar interneurons during air-stepping in subchronic spinal cats. *J Neurophysiol* **105**, 1011–1022, DOI: 10.1152/jn.00523.2010. (2011).

**23.** Bauer, U. Ripser: efficient computation of Vietoris–Rips persistence barcodes. *J. Appl. Comput. Topol.* **5**, 391–423, DOI: 10.1007/s41468-021-00071-5 (2021).

**24.** Perich, M. G., Narain, D. & Gallego, J. A. A neural manifold view of the brain. *Nat. Neurosci.* DOI: 10.1038/s41593-025-02031-z (2025).

**25.** Mitchell-Heggs, R., Prado, S., Gava, G. P., Go, M. A. & Schultz, S. R. Neural manifold analysis of brain circuit dynamics in health and disease. *J. Comput. Neurosci.* **51**, 1–21, DOI: 10.1007/s10827-022-00839-3 (2023).

**26.** Gardner, R. J. *et al.* Toroidal topology of population activity in grid cells. *Nature* **602**, 123–128, DOI: 10.1038/s41586-021-04268-7 (2022).

**27.** Martiny, E. S., Jensen, M. H. & Heltberg, M. S. Detecting limit cycles in stochastic time series. *Phys. A: Stat. Mech. its Appl.* **605**, 127917, DOI: 10.1016/j.physa.2022.127917 (2022).

**28.** Kim, S. S., Rouault, H., Druckmann, S. & Jayaraman, V. Ring attractor dynamics in the Drosophila central brain. *Science* **356**, 849–853, DOI: 10.1126/science.aal4835 (2017).

**29.** Khona, M. & Fiete, I. R. Attractor and integrator networks in the brain. *Nat. Rev. Neurosci.* **23**, 744–766, DOI: 10.1038/s41583-022-00642-0 (2022).

**30.** Voroslakos, M. *et al.* 3D-printed Recoverable Microdrive and Base Plate System for Rodent Electrophysiology. *BIO-PROTOCOL* **11**, DOI: 10.21769/BioProtoc.4137 (2021).

**31.** Chettih, S. N. & Harvey, C. D. Single-neuron perturbations reveal feature-specific competition in V1. *Nature* **567**, 334–340, DOI: 10.1038/s41586-019-0997-6 (2019).

**32.** Kaur, J. & Berg, R. W. Viral strategies for targeting spinal neuronal subtypes in adult wild-type rodents. *Sci. Reports* **12**, 8627, DOI: 10.1038/s41598-022-12535-4 (2022).

**33.** Jun, J. J. *et al.* Fully integrated silicon probes for high-density recording of neural activity. *Nature* **551**, 232–236, DOI: 10.1038/nature24636 (2017).

**34.** Berg, R. W., Chen, M.-T., Huang, H.-C., Hsiao, M.-C. & Cheng, H. A method for unit recording in the lumbar spinal cord during locomotion of the conscious adult rat. *J. Neurosci. Methods* **182**, DOI: 10.1016/j.jneumeth.2009.05.023 (2009).

**35.** Kaur, J. *et al.* Pedunculopontine-stimulation obstructs hippocampal theta rhythm and halts movement. *Sci. Reports* **15**, 17903, DOI: 10.1038/s41598-025-01695-8 (2025).

**36.** Avants, B. B. *et al.* A reproducible evaluation of ANTs similarity metric performance in brain image registration. *NeuroImage* **54**, 2033–2044, DOI: 10.1016/j.neuroimage.2010.09.025 (2011).

**37.** Pachitariu, M., Sridhar, S., Pennington, J. & Stringer, C. Spike sorting with Kilosort4. *Nat. Methods* **21**, 914–921, DOI: 10.1038/s41592-024-02232-7 (2024).

**38.** Mathis, A. *et al.* DeepLabCut: markerless pose estimation of user-defined body parts with deep learning. *Nat. Neurosci.* **21**, 1281–1289, DOI: 10.1038/s41593-018-0209-y (2018).

**39.** Bácskai, T., Rusznák, Z., Paxinos, G. & Watson, C. Musculotopic organization of the motor neurons supplying the mouse hindlimb muscles: a quantitative study using Fluoro-Gold retrograde tracing. *Brain Struct. Funct.* **219**, 303–321, DOI: 10.1007/s00429-012-0501-7 (2014).

**40.** Veshchitskii, A., Shkorbatova, P. & Merkulyeva, N. Distribution of the motoneuronal pools controlling the hindlimb muscles in the lumbar spinal cord of the Acomys cahirinus. *Brain Struct. Funct.* **230**, 86, DOI: 10.1007/s00429-025-02946-0 (2025).

**41.** Paxinos, G. & Watson, C. *The Rat Brain in Stereotaxic Coordinates : Hard Cover Edition* (Elsevier Science, 2013).

**42.** Molander, C., Xu, Q. & Grant, G. The cytoarchitectonic organization of the spinal cord in the rat. I. The lower thoracic and lumbosacral cord. *J. Comp. Neurol.* **230**, 133–141, DOI: 10.1002/cne.902300112 (1984).
